# A one-enzyme RT-qPCR assay for SARS-CoV-2, and procedures for reagent production

**DOI:** 10.1101/2020.03.29.013342

**Authors:** Sanchita Bhadra, Andre C. Maranhao, Andrew D. Ellington

## Abstract

Given the scale of the ongoing COVID-19 pandemic, the need for reliable, scalable testing, and the likelihood of reagent shortages, especially in resource-poor settings, we have developed a RT-qPCR assay that relies on an alternative to conventional viral reverse transcriptases, a thermostable reverse transcriptase / DNA polymerase (RTX)^1^. Here we show that RTX performs comparably to the other assays sanctioned by the CDC and validated in kit format. We demonstrate two modes of RTX use – (i) dye-based RT-qPCR assays that require only RTX polymerase, and (ii) TaqMan RT-qPCR assays that use a combination of RTX and Taq DNA polymerases (as the RTX exonuclease does not degrade a TaqMan probe). We also provide straightforward recipes for the purification of this alternative reagent. We anticipate that in low resource or point-of-need settings researchers could obtain the available constructs from Addgene or our lab and begin to develop their own assays, within whatever regulatory framework exists for them.

We lay out protocols for dye-based and TaqMan probe-based assays, in order to best compare with ‘gold standard’ reagents. These protocols should form the basis of further modifications that can simplify the assay to the use of overexpressing cells themselves as reagents.

<u>Developing dye-based and TaqMan probe-based RT-qPCR assays with RTX</u>

## MATERIALS

- 10X RTX buffer: 600 mM Tris-HCl, 250 mM (NH_4_)_2_SO_4_, 100 mM KCl, 20 mM MgSO_4_, pH 8.4 @25°C
- 10X ThermoPol buffer (New England Biolabs): 200 mM Tris-HCl, 100 mM (NH_4_)_2_SO_4_, 100 mM KCl, 20 mM MgSO_4_, 1% Triton^®^ X-100, pH 8.8 @25°C
- Forward PCR primer^a^
- Reverse PCR primer^a^
- TaqMan probe^a^
- 20X EvaGreen solution
- 4 mM deoxyribonucleotides (dNTP)
- 5M Betaine
- 20 U/µL SUPERase.In RNase inhibitor
- 5 U/µL Taq DNA polymerase
- 0.2 mg/ml RTX polymerase^b^
- Nuclease-free water
- qPCR tubes or plates^c^
- Real-time PCR machine^c^
- Cold-block or ice
- Barrier tips

a. Individual primer and probe stocks or pre-made primer-probe mixes for CDC N1, N2 and N3 assays (**Table 1**) available from Integrated DNA Technologies may be used.
b. RTX polymerase with proofreading capability (RTX), and RTX polymerase without proofreading capability (RTX Exo-) have been compared.
c. LightCycler96 real-time PCR machine and 96-well plates with optical plastic film cover designed for use with LightCycler platform were used for these experiments.

**Table 1.**
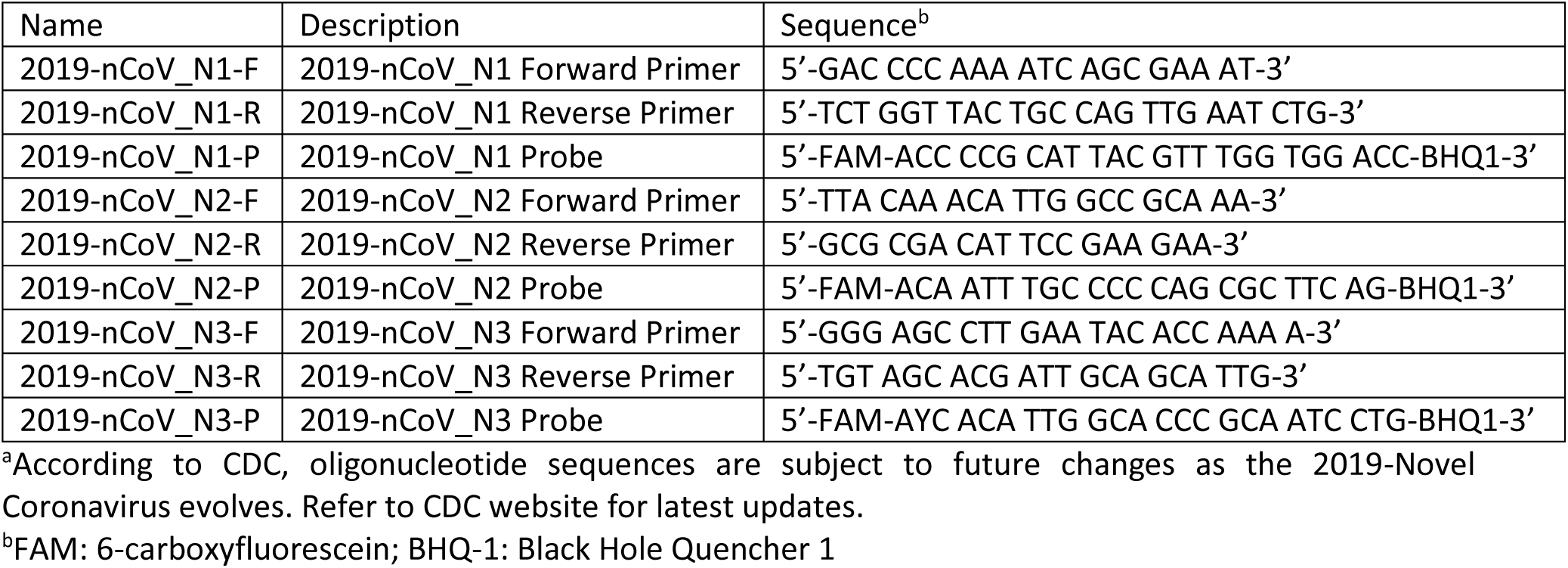
CDC TaqMan qRT-PCR primers and probes for SARS-CoV-2a (adapted from https://www.cdc.gov/coronavirus/2019-ncov/lab/rt-pcr-panel-primer-probes.html).

## RESULTS

### Dye-based RT-qPCR reaction set up using only RTX DNA polymerase

EvaGreen dye-containing RT-qPCR reaction set up shown in **Table 2** was used for assembling CDC SARS-CoV-2 N1, N2, and N3 assays. The three assays target three different regions within the nucleocapsid phosphoprotein (N) gene near the 3’-end of the ∼30 kb viral genome. In particular, N1 primers (**Table 1**) amplify a 72 nt long region from position 28287 to 28358 of the genome (GenBank Accession No. MN985325.1). The N2 primers produce a 67 bp amplicon from the genomic position 29164 to 29230. Meanwhile, the N3 primers amplify a 72 nt long region from genomic position 28681 to 28752. All reaction mixes were assembled on a cold-block and directly transferred to a real-time PCR machine programmed to cycle through the steps depicted in **Table 3**. Melt curve analysis was included after PCR amplification in order to distinguish target amplicons from spurious products.

**Table 2.** EvaGreen RT-qPCR reaction mix using only RTX DNA polymerase

|  | Volume (μL) | Final concentration |
| --- | --- | --- |
| RNA sample | 5 |  |
| Forward PCR primer | 1.5 | 500 nM |
| Reverse PCR primer | 1.5 | 500 nM |
| 20X EvaGreen | 1.25 | 1X |
| 10X RTX buffer | 2.5 | 1X |
| 4 mM dNTP | 2.5 | 400 μM |
| 20 U/μL SUPERase.In RNase inhibitor | 1 | 0.8 U/μL |
| RTX or RTX Exo- 0.2mg/ml | 0.5 | 4 ng/μL |
| 5M betaine | 2.5 | 0.5M |
| Water | 6.75 |  |
| Total reaction volume | 25 |  |

**Table 3.** RTX polymerase-based EvaGreen RT-qPCR cycling conditions

| Step | Temperature | Duration | Signal acquisition |
| --- | --- | --- | --- |
| 1<sup>a</sup> | 60 °C | 30 min | None |
| 2<sup>a</sup> | 95 °C | 10 min | None |
| 55 cycles of Steps 3 and 4 |  |  |  |
| 3 | 95 °C | 15 sec | None |
| 4<sup>b</sup> | 55 °C | 30 sec | Single |
| Melt curve analysis in Steps 5, 6, and 7 |  |  |  |
| 5 | 95 °C | 10 sec | None |
| 6 | 65 °C | 60 sec | None |
| 7 | 97 °C | 1 sec | Continuous<sup>c</sup> |

<sup>a</sup>Assay time may be reduced by shortening reverse transcription step (Step 1) to 15 min and the following denaturation step (Step 2) to 2 min.
<sup>b</sup>Signal acquisition occurs in the FAM (fluorescein) channel during step 4.
<sup>c</sup>20 measurements/°C temperature change

Representative results of SARS-CoV-2 RT-qPCR tests performed using synthetic RNA templates and only RTX or RTX Exo-DNA polymerases are depicted in **Figures 1 and 2**.

**Figure 1.**
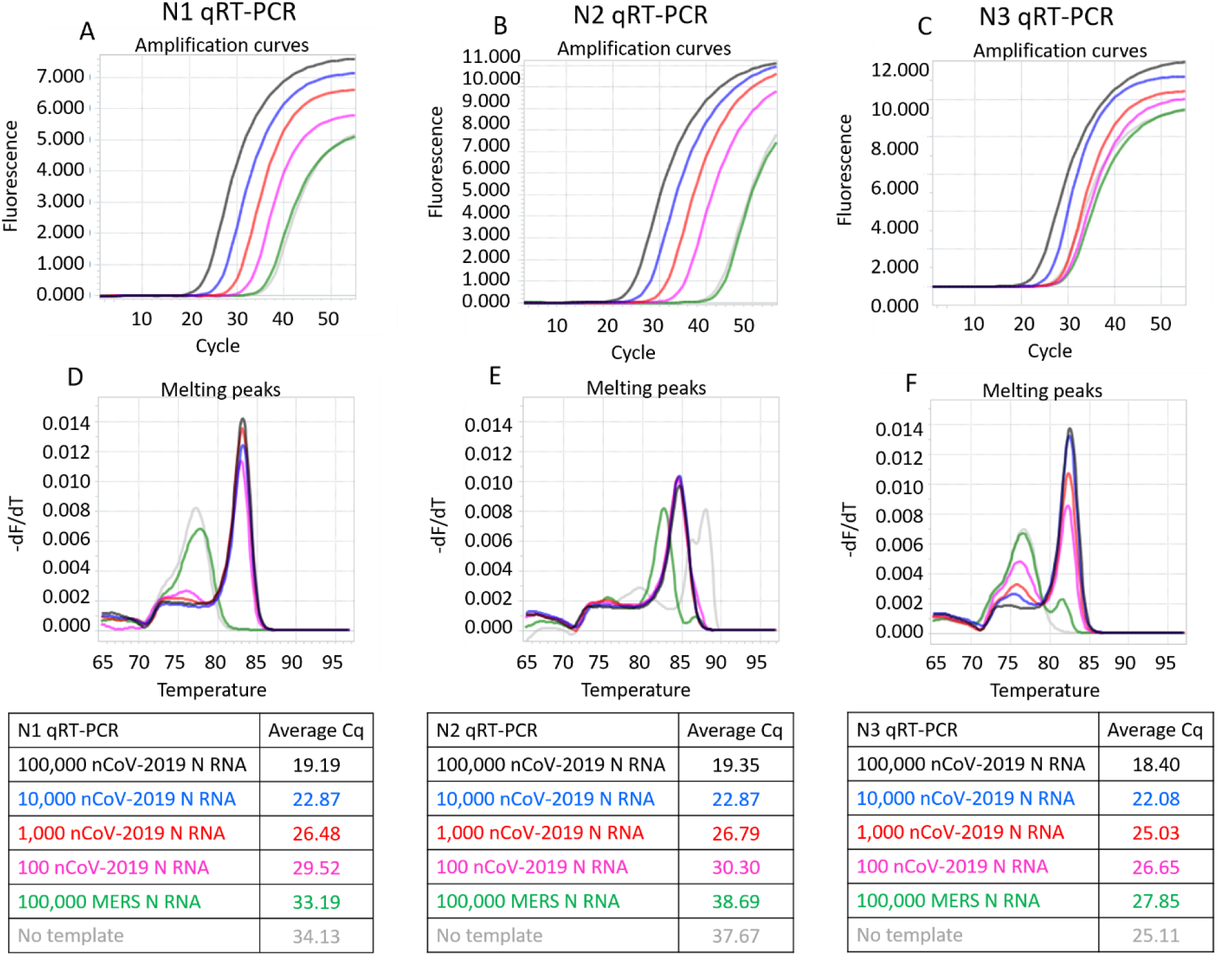
CDC SARS-CoV-2 N1, N2, and N3 RT-qPCR assays performed using indicated copies of synthetic RNA and RTX DNA polymerase. Panels A, B, and C depict N1, N2, and N3 assays measured in real-time using EvaGreen dye. Amplification curves from reactions containing 100,000 (black traces), 10,000 (blue traces), 1,000 (red traces), and 100 (pink traces) copies of SARS-CoV-2 synthetic N RNA are depicted. Negative control reactions either contained no templates (gray traces) or contained 100,000 copies of synthetic N RNA from MERS-CoV (green traces). Panels D, E, and F depict melting peaks of amplicons determined using the ‘Tm calling’ analysis in the LightCycler96 software. Average Cq values of all assays are tabulated.

**Figure 2.**
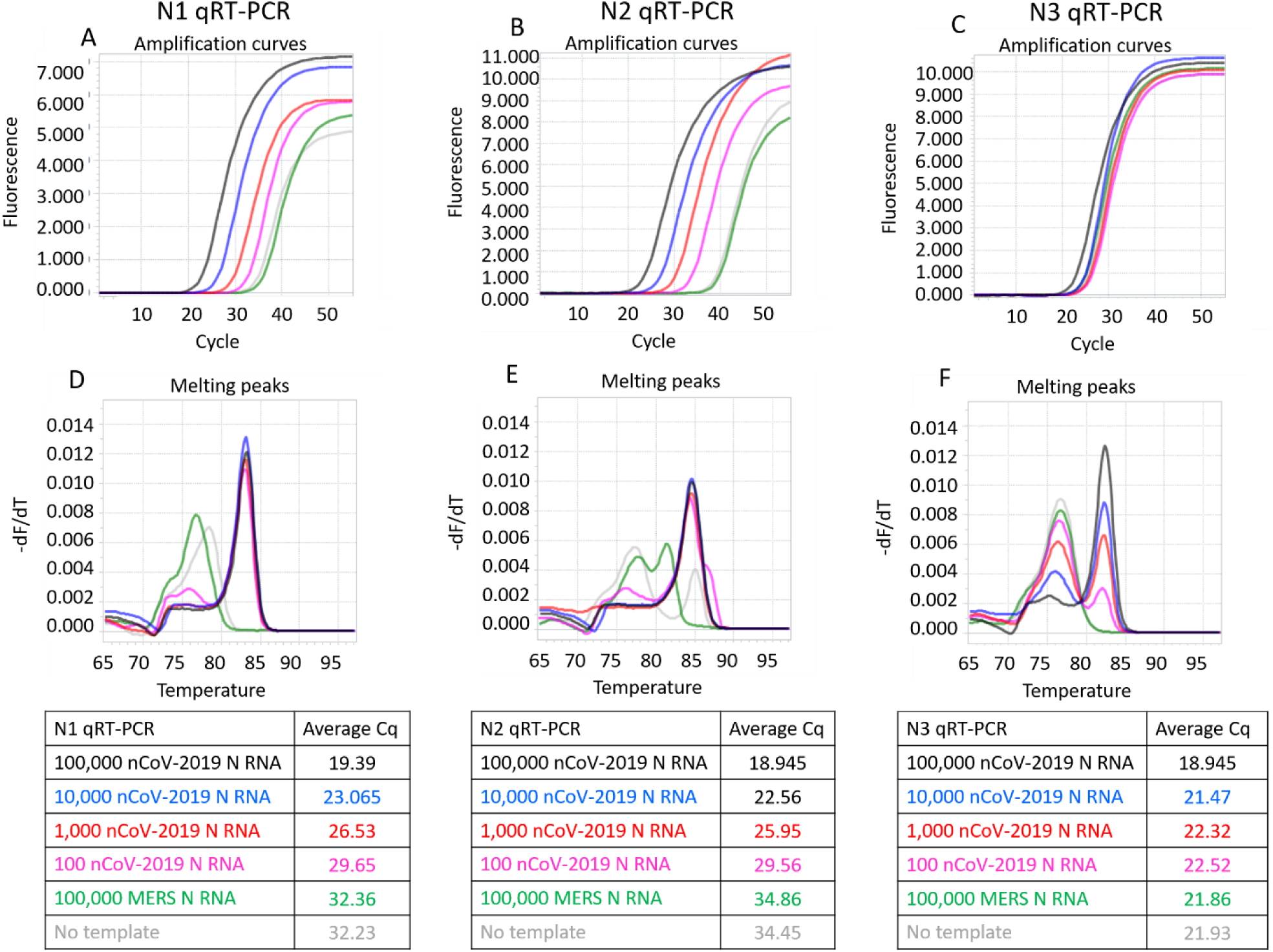
CDC SARS-CoV-2 N1, N2, and N3 RT-qPCR assays performed using indicated copies of synthetic RNA and RTX Exo-DNA polymerase. Panels A, B, and C depict N1, N2, and N3 assays measured in real-time using EvaGreen dye. Amplification curves from reactions containing 100,000 (black traces), 10,000 (blue traces), 1,000 (red traces), and 100 (pink traces) copies of SARS-CoV-2 synthetic N RNA are depicted. Negative control reactions either contained no templates (gray traces) or contained 100,000 copies of synthetic N RNA from MERS-CoV (green traces). Panels D, E, and F depict melting peaks of amplicons determined using the ‘Tm calling’ analysis in the LightCycler96 software. Average Cq values of all assays are tabulated.

These results demonstrate that RTX DNA polymerase alone, with or without proofreading capability, can support dye-based RT-qPCR analyses. In our hands the full-length RTX DNA polymerase was slightly better than the Exo-version, especially in the N3 assay.

### RTX polymerase-based TaqMan qRT-PCR reaction set up

In order to perform TaqMan RT-qPCR we developed a one-pot two enzyme system comprising RTX and Taq DNA polymerases. TaqMan qRT-PCR reaction set up shown in **Table 4** was used for assembling CDC SARS-CoV-2 N1, N2, and N3 assays using either RTX buffer or ThermoPol buffer. Control reactions lacking RTX polymerase were set up using the same recipe except RTX was replaced with water. All reaction mixes were assembled on a cold-block and directly transferred to a real-time PCR machine programmed to cycle through the steps depicted in **Table 5**.

**Table 4.**
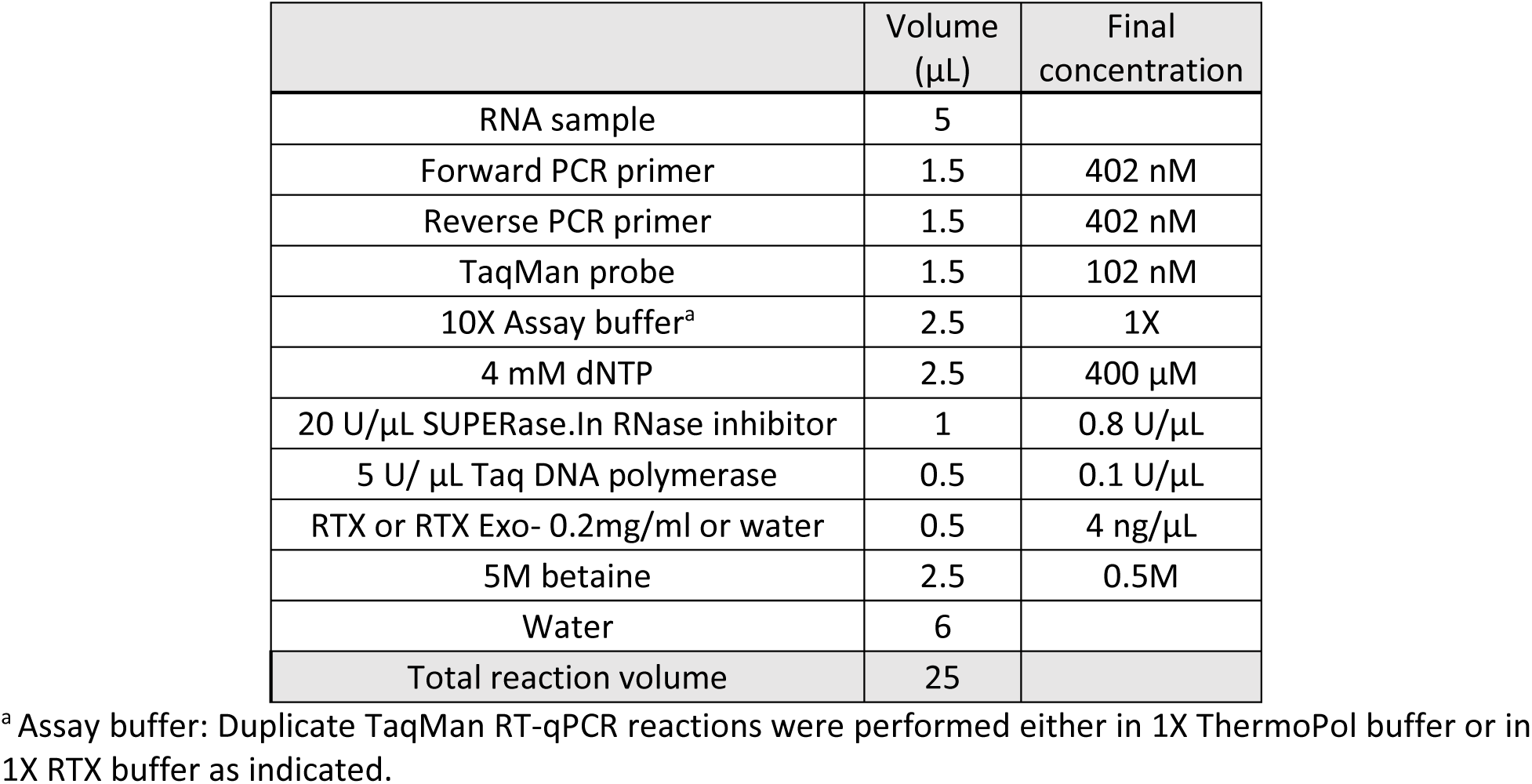
RTX polymerase-based TaqMan qRT-PCR reaction mix

|  | Volume (μL) | Final concentration |
| --- | --- | --- |
| RNA sample | 5 |  |
| Forward PCR primer | 1.5 | 402 nM |
| Reverse PCR primer | 1.5 | 402 nM |
| TaqMan probe | 1.5 | 102 nM |
| 10X Assay buffer<sup>a</sup> | 2.5 | 1X |
| 4 mM dNTP | 2.5 | 400 μM |
| 20 U/μL SUPERase.In RNase inhibitor | 1 | 0.8 U/μL |
| 5 U/ μL Taq DNA polymerase | 0.5 | 0.1 U/μL |
| RTX or RTX Exo- 0.2mg/ml or water | 0.5 | 4 ng/μL |
| 5M betaine | 2.5 | 0.5M |
| Water | 6 |  |
| Total reaction volume | 25 |  |

<sup>a</sup> Assay buffer: Duplicate TaqMan RT-qPCR reactions were performed either in 1X ThermoPol buffer or in 1X RTX buffer as indicated.

**Table 5.** RTX polymerase-based TaqMan qRT-PCR cycling conditions

|  |  |  |
| --- | --- | --- |
| Step 1<sup>a</sup> | 60 °C | 30 min |
| Step 2<sup>a</sup> | 95 °C | 10 min |
| 55 cycles of Steps 3 and 4 |  |  |
| Step 3 | 95 °C | 15 sec |
| Step 4<sup>b</sup> | 55 °C | 30 sec |

<sup>a</sup> Assay time may be reduced by shortening reverse transcription step (Step 1) to 15 min and the following denaturation step (Step 2) to 2 min.
<sup>b</sup> Signal acquisition occurs in the FAM (fluorescein) channel during step 4.

Representative results of RTX polymerase-based SARS-CoV-2 TaqMan qRT-PCR tests performed using synthetic RNA templates are depicted in **Figures 3 and 4**. Both RTX and RTX Exo-when combined with Taq DNA polymerase were able to support one-pot TaqMan RT-qPCR assays for all three CDC SARS-CoV-2 assays. Under both buffer conditions tested, 1X ThermoPol versus 1X RTX buffer, RTX polymerase yielded more consistent amplification curves for all three CDC assays compared to RTX Exo-polymerase.

**Figure 3.**
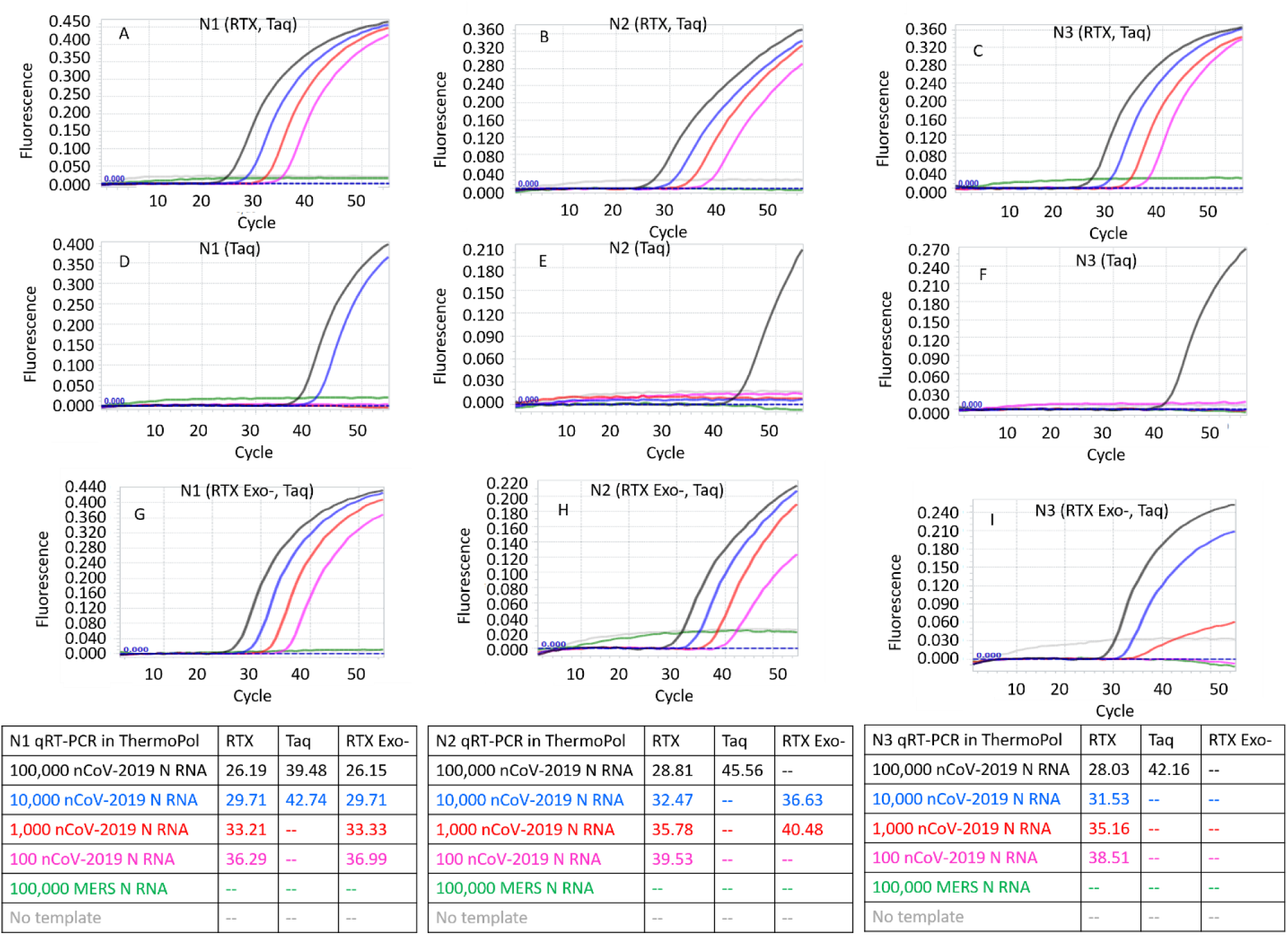
CDC SARS-CoV-2 N1, N2, and N3 TaqMan RT-qPCR assays performed in 1X ThermoPol buffer using indicated copies of synthetic RNA and RTX or RTX Exo- and Taq DNA polymerases. Panels A, B, and C depict TaqMan assays containing both RTX and Taq DNA polymerases. Panels D, E, and F depict TaqMan assays containing only Taq DNA polymerase. Panels G, H, and I depict TaqMan assays containing RTX Exo- and Taq DNA polymerases. Amplification curves from reactions containing 100,000 (black traces), 10,000 (blue traces), 1,000 (red traces), and 100 (pink traces) copies of SARS-CoV-2 synthetic N RNA are depicted. Negative control reactions either contained no templates (gray traces) or contained 100,000 copies of synthetic N RNA from MERS-CoV (green traces). Cq values of all assays are tabulated.

**Figure 4.**
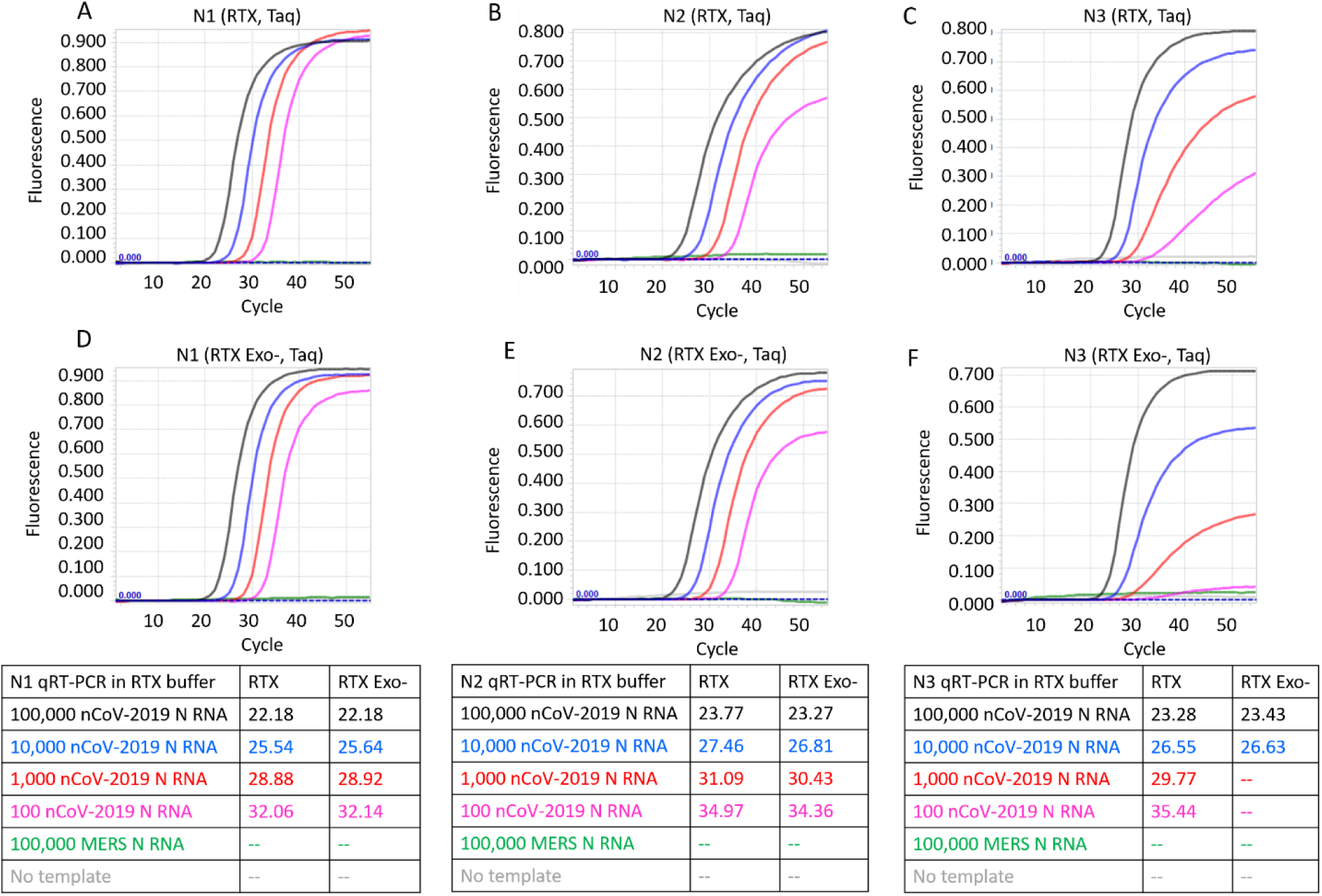
CDC SARS-CoV-2 N1, N2, and N3 TaqMan RT-qPCR assays performed in 1X RTX buffer using indicated copies of synthetic RNA and RTX or RTX Exo- and Taq DNA polymerases. Panels A, B, and C depict TaqMan assays containing RTX and Taq DNA polymerases. Panels D, E, and F depict TaqMan assays containing RTX Exo- and Taq DNA polymerases. Amplification curves from reactions containing 100,000 (black traces), 10,000 (blue traces), 1,000 (red traces), and 100 (pink traces) copies of SARS-CoV-2 synthetic N RNA are depicted. Negative control reactions either contained no templates (gray traces) or contained 100,000 copies of synthetic N RNA from MERS-CoV (green traces). Cq values of all assays are tabulated.

To compare the RTX-based TaqMan qRT-PCR with qRT-PCR performed using commercial master mix, duplicate reactions (**Table 6**) were set up using TaqPath™ 1-Step RT-qPCR Master Mix, CG (Thermo Fisher Scientific, Waltham, MA, USA). All reactions were assembled on cold block and then incubated at room temperature (25 °C benchtop) for 2 min prior to being loaded on a real-time PCR machine set to cycle through the steps indicated in **Table 7**.

**Table 6.**
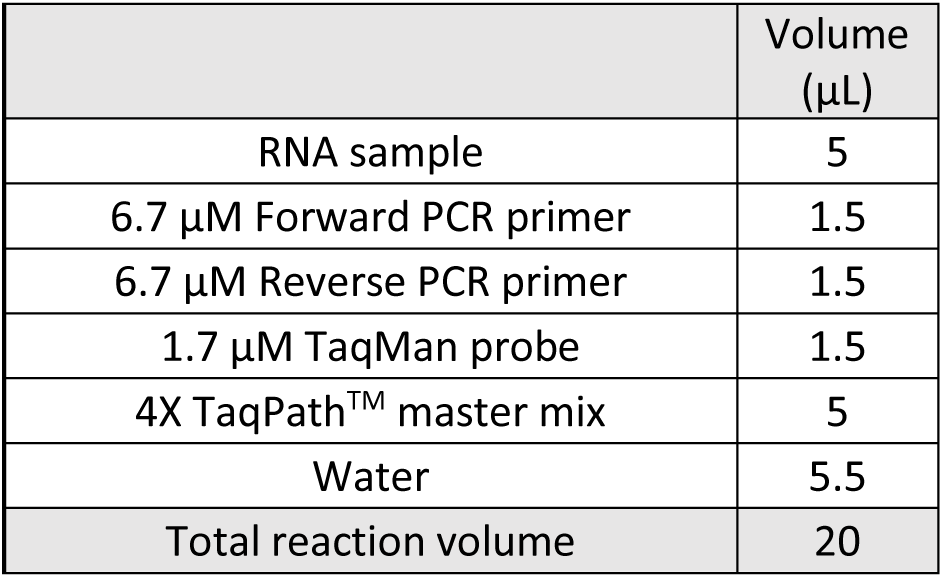
CDC N1, N2, N3 TaqMan qRT-PCR assay set up using commercial qRT-PCR master mix

**Table 7.** TaqPath™ TaqMan qRT-PCR cycling conditions

|  |  |  |
| --- | --- | --- |
| Step 1 | 50 °C | 15 min |
| Step 2 | 95 °C | 2 min |
| 45 cycles of Steps 3 and 4 |  |  |
| Step 3 | 95 °C | 3 sec |
| Step 4 | 55 °C | 30 sec |

**Figure 5** depicts typical results from CDC N1, N2, and N3 TaqMan qRT-PCR reactions performed using synthetic RNA and TaqPath™ commercial qRT-PCR mastermix.

**Figure 5.**
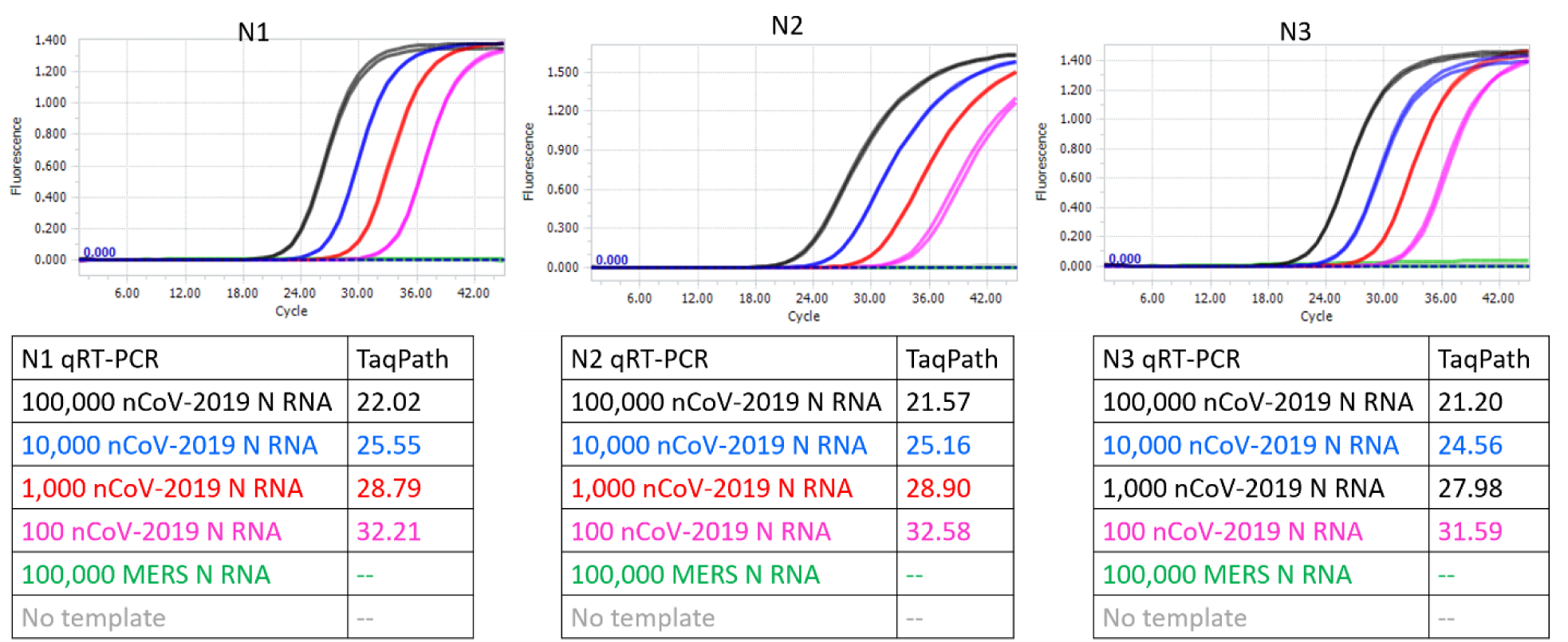
CDC SARS-CoV-2 N1, N2, and N3 TaqMan qRT-PCR assays performed using indicated copies of synthetic RNA and TaqPath™ commercial qRT-PCR mastermix. Amplification curves from reactions containing 100,000 (black traces), 10,000 (blue traces), 1,000 (red traces), and 100 (pink traces) copies of SARS-CoV-2 synthetic N RNA are depicted. Negative control reactions either contained no templates (gray traces) or contained 100,000 copies of synthetic N RNA from MERS-CoV (green traces). Cq values of all assays are tabulated.

The results with either a viral RT (in TaqPath™ commercial mastermix) or RTX are summarized in **Table 8**. As can be seen, RTX-based TaqMan assays are of comparable sensitivity and specificity to the gold standard TaqPath assay. While all three RTX-based TaqMan assays performed in ThermoPol buffer consistently yielded Cq values, these were somewhat delayed compared to Cq values obtained with the commercial TaqPath™ Master Mix. In contrast, RTX-based TaqMan assays when executed in RTX buffer yielded Cq values that were closer to Cq values obtained with the commercial mastermix.

**Table 8.**
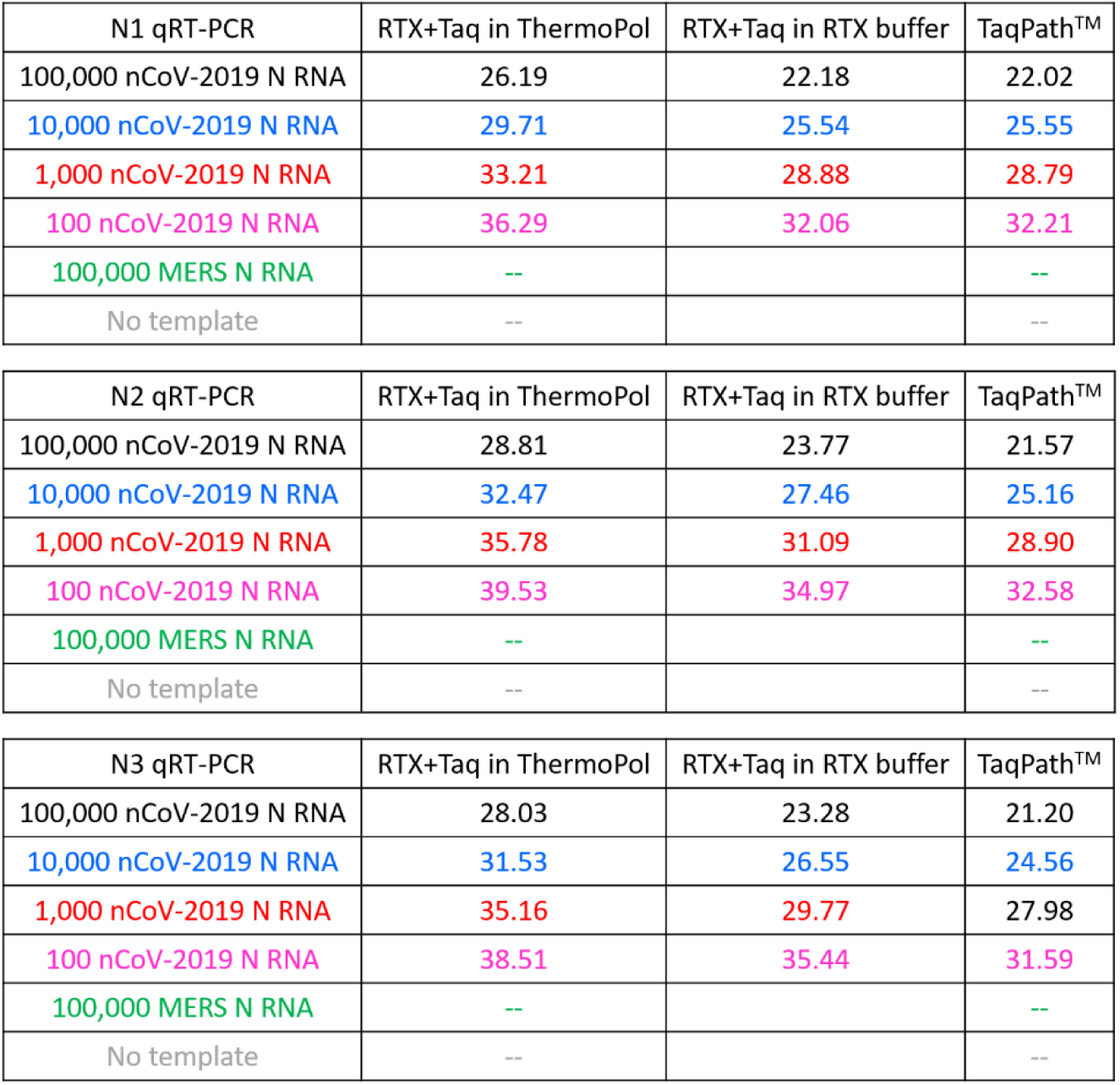
Comparison of Cq values of RTX-based and TaqPath-based SARS-CoV-2 TaqMan qRT-PCR assays

| N1 qRT-PCR | RTX+Taq in ThermoPol | RTX+Taq in RTX buffer | TaqPath™ |
| --- | --- | --- | --- |
| 100,000 nCoV-2019 N RNA | 26.19 | 22.18 | 22.02 |
| 10,000 nCoV-2019 N RNA | 29.71 | 25.54 | 25.55 |
| 1,000 nCoV-2019 N RNA | 33.21 | 28.88 | 28.79 |
| 100 nCoV-2019 N RNA | 36.29 | 32.06 | 32.21 |
| 100,000 MERS N RNA | -- |  | -- |
| No template | -- |  | -- |

| N2 qRT-PCR | RTX+Taq in ThermoPol | RTX+Taq in RTX buffer | TaqPath™ |
| --- | --- | --- | --- |
| 100,000 nCoV-2019 N RNA | 28.81 | 23.77 | 21.57 |
| 10,000 nCoV-2019 N RNA | 32.47 | 27.46 | 25.16 |
| 1,000 nCoV-2019 N RNA | 35.78 | 31.09 | 28.90 |
| 100 nCoV-2019 N RNA | 39.53 | 34.97 | 32.58 |
| 100,000 MERS N RNA | -- |  | -- |
| No template | -- |  | -- |

| N3 qRT-PCR | RTX+Taq in ThermoPol | RTX+Taq in RTX buffer | TaqPath™ |
| --- | --- | --- | --- |
| 100,000 nCoV-2019 N RNA | 28.03 | 23.28 | 21.20 |
| 10,000 nCoV-2019 N RNA | 31.53 | 26.55 | 24.56 |
| 1,000 nCoV-2019 N RNA | 35.16 | 29.77 | 27.98 |
| 100 nCoV-2019 N RNA | 38.51 | 35.44 | 31.59 |
| 100,000 MERS N RNA | -- |  | -- |
| No template | -- |  | -- |

#### Summary of assay performance

Overall, RTX performs well either as a single enzyme for RT-qPCR with intercalating dyes, such as EvaGreen, or as the RT component of a TaqMan based assay. At the lowest RNA concentrations examined (100 copies), the gold standard TaqPath assay general gave a signal at a Cq value of ca. 32. The more robust full-length RTX on its own actually performed slightly better, with C_q_ values typically from 29-30 (although the N3 primer set gives higher signal and higher background). When used as a substitute for a viral RT, RTX is slightly less sensitive overall, with Cq values typically from 32-35.

#### Purification of RTX

For sites wishing to produce their own RTX, the following protocol can be used.

**Table 7.**
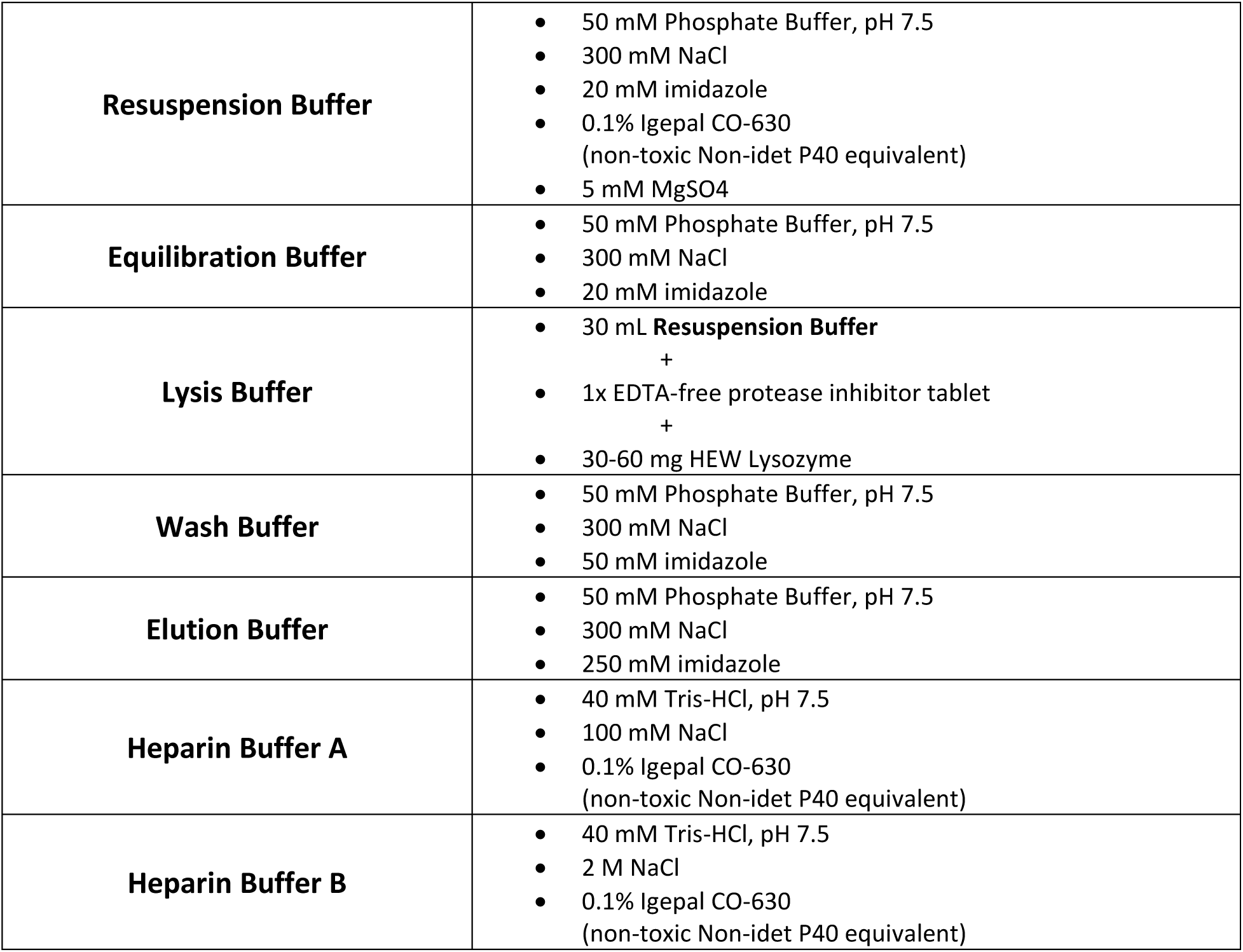
Buffers for RTX purification

|  |  |
| --- | --- |
| **Resuspension Buffer** | • 50 mM Phosphate Buffer, pH 7.5<br>• 300 mM NaCl<br>• 20 mM imidazole<br>• 0.1% Igepal CO-630 (non-toxic Non-idet P40 equivalent)<br>• 5 mM MgSO<sub>4</sub> |
| **Equilibration Buffer** | • 50 mM Phosphate Buffer, pH 7.5<br>• 300 mM NaCl<br>• 20 mM imidazole |
| **Lysis Buffer** | • 30 mL **Resuspension Buffer**<br>• +<br>• 1x EDTA-free protease inhibitor tablet<br>• +<br>• 30-60 mg HEW Lysozyme |
| **Wash Buffer** | • 50 mM Phosphate Buffer, pH 7.5<br>• 300 mM NaCl<br>• 50 mM imidazole |
| **Elution Buffer** | • 50 mM Phosphate Buffer, pH 7.5<br>• 300 mM NaCl<br>• 250 mM imidazole |
| **Heparin Buffer A** | • 40 mM Tris-HCl, pH 7.5<br>• 100 mM NaCl<br>• 0.1% Igepal CO-630 (non-toxic Non-idet P40 equivalent) |
| **Heparin Buffer B** | • 40 mM Tris-HCl, pH 7.5<br>• 2 M NaCl<br>• 0.1% Igepal CO-630 (non-toxic Non-idet P40 equivalent) |

**Table 8.**
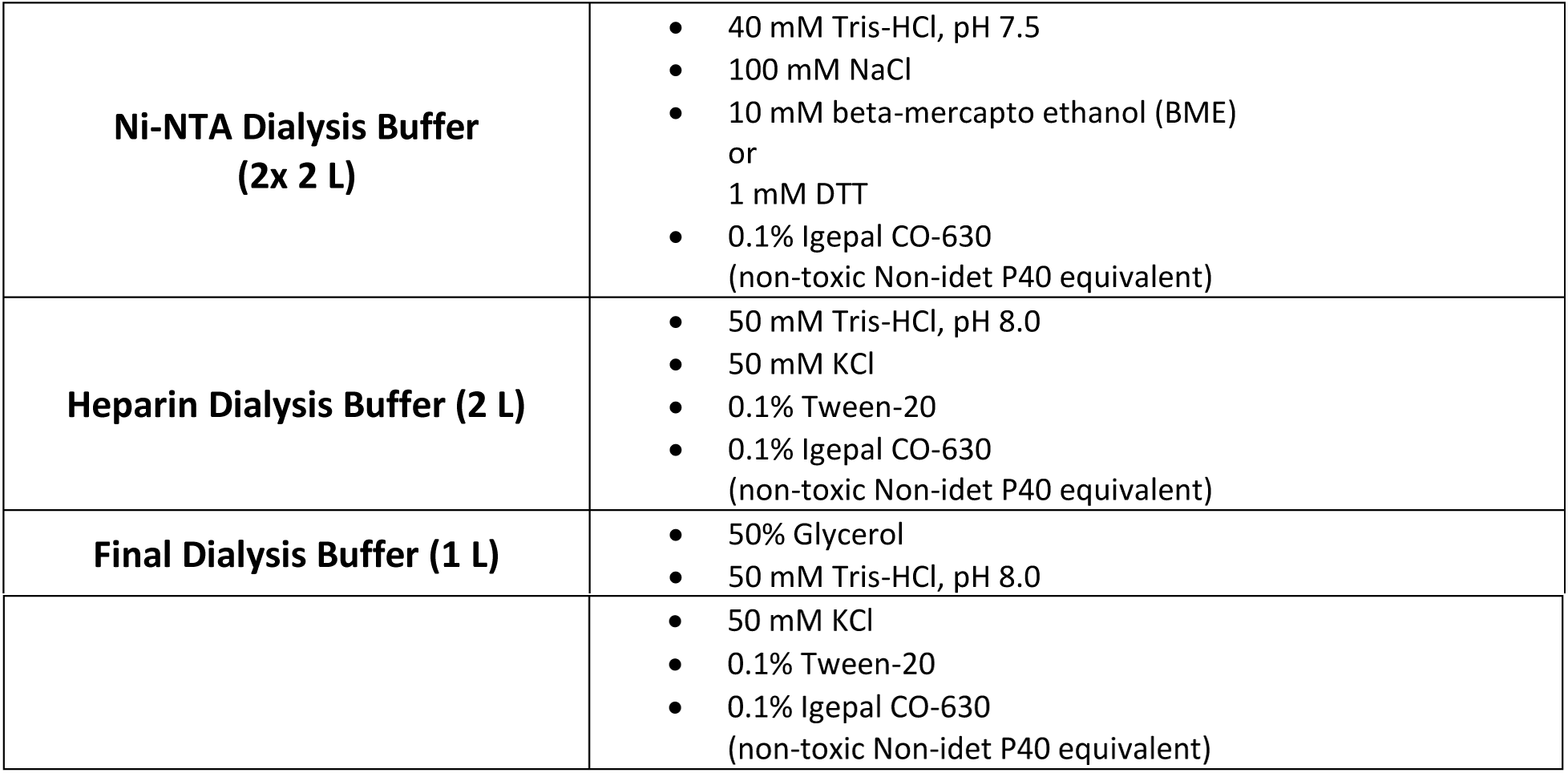
Buffers for dialysis of RTX

In brief, a T7-based *E. coli* expression strain such as BL21(DE3) was transformed with a RTX expression plasmid that contained an carboxyl terminal six histidine tag for purification. On the next day, an individual transformant colony was picked into desired media with appropriate antibiotics (e.g. ampicillin) and 1-2% glucose. This starter culture was grown overnight in shaking incubator at 37°C.

On day three, 1 L Superior Broth™ (Athena Enzyme Systems™) or other rich media with appropriate antibiotics in a 4 L Erlenmeyer flask was inoculated with 250-350 μL of the overnight starter culture. Incubate at 37°C, 250 rpm until the culture reaches OD_600_ of 0.4-0.7 (0.5-0.6 is ideal). When desired OD_600_ is reached, place flask(s) in 4°C cold room for 30-45 min and set shaking incubator to 18°C. Following cold incubation, induce expression of RTX by adding 1 mL of 1 M IPTG to each 1 L of culture. Place flask(s) in the 18°C shaking incubator for 16-18 hours.

Following overnight expression, cells were harvested by centrifuging cultures at 4°C, 5000 xg for 20 min. On ice, resuspend cell pellet in 30 mL cold **Lysis Buffer**. Transfer the resuspended cell pellet to a small 50 mL beaker with clean stir bar and place securely in an ice bath. With moderate stirring, sonicate the sample using 40% amplitude and 1 sec ON / 4 sec OFF for 4 min total sonication time. Centrifuge the resulting lysate at 4°C, 35,000 xg for 30 min. Carefully transfer the supernatant to a clean ultracentrifugation tube, shake in thermomixer at 400 rpm, 85°C for 10 min, and then place on ice for 10 min. Centrifuged the heat-treated lysate at 4°C, 35,000 xg for 30 min. Carefully transfer the supernatant to a clean tube and filter the clarified lysate using a 0.2 μm filter.

Perform IMAC column purification in a cold room or allow gravity flow to occur in a refrigerator. Prepare Ni-NTA agarose columns with final column volume (CV) of 1 mL. Apply the clarified lysate from 1 L of expressed culture to a previously equilibrated column and collect the flow-through. Wash column with 20 CV **Equilibration Buffer** and collect the flow-through. Wash column with 5 CV **Wash Buffer** and collect flow-through. Elute RTX from column with 5 mL **Elution Buffer** and transfer to a dialysis cassette with appropriate molecular weight cut-off. Dialyze eluate into 2 L of **Ni-NTA Dialysis Buffer** for 3-4 hours at 4°C. Then dialyze eluate into a second 2 L of **Ni-NTA Dialysis Buffer** overnight at 4°C.

Dialyzed eluate was then passed over an equilibrated 5 mL heparin column (HiTrap™ Heparin HP) and eluted along a sodium chloride gradient (100 mM to 2 M NaCl). The peak corresponding to RTX or its variants could be expected between 40-60% Buffer B. RTX fractions were collected, pooled, and dialyzed into 2 L **Heparin Dialysis Buffer** for 3-4 hours. Then dialysis cassettes were transferred into 2 L **Final Dialysis Buffer** overnight where it is expected that the protein sample volume will decrease significantly. Following the final dialysis, protein sample was recovered and protein concentration was determined before storing at −20°C.

A similar protocol should work for Taq and other thermostable polymerases, but may require different dialysis and storage buffers.

#### Summary and opportunities for distribution

By obtaining a RTX expression vector and purifying the thermostable reverse transcriptase it should prove possible to carry out RT-qPCR reactions with a sensitivity similar to that already observed for approved kits, with few or no false positive results. The sequence information for plasmids for RTX [pET_RTX-6xHis] and its exonuclease deficient variant, RTX(exo-) [pET_RTX(exo -)-6xHis], bearing a C-terminal 6xHis tags can be found in the **Supplemental Information**. These plasmids will be uploaded to Addgene in short order, or can be obtained via https://reclone.org/. The Board of Regents of The University of Texas has licensed IP covering RTX to Promega Corporation.

## Acknowledgements

We acknowledge the extraordinary contributions of the Schoggins lab in making synthetic RNA available on short notice for these assays.

## Supporting Information

**Figure S1.**
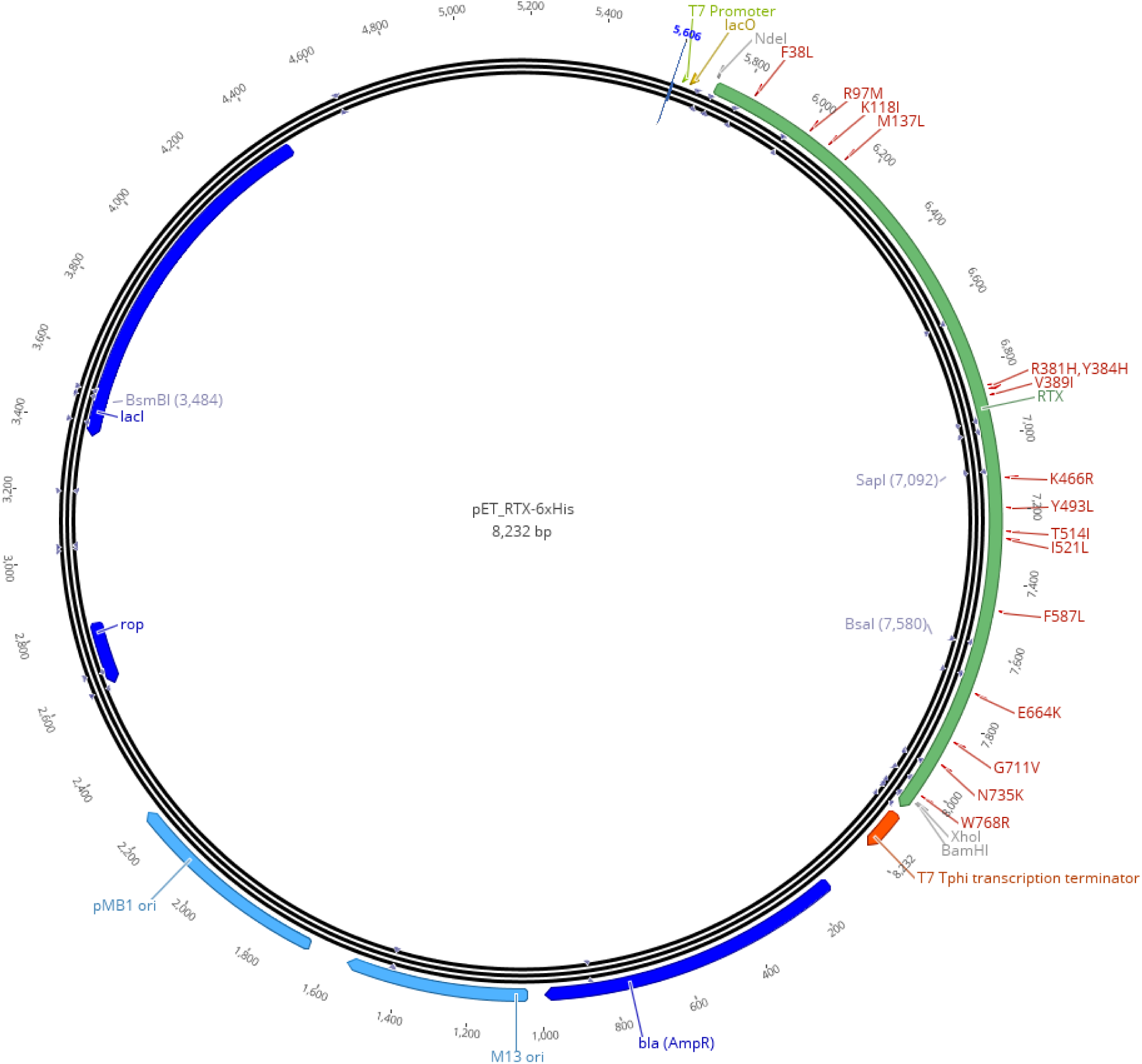
Plasmid map of pET_RTX-6xHis

**Table S1.**
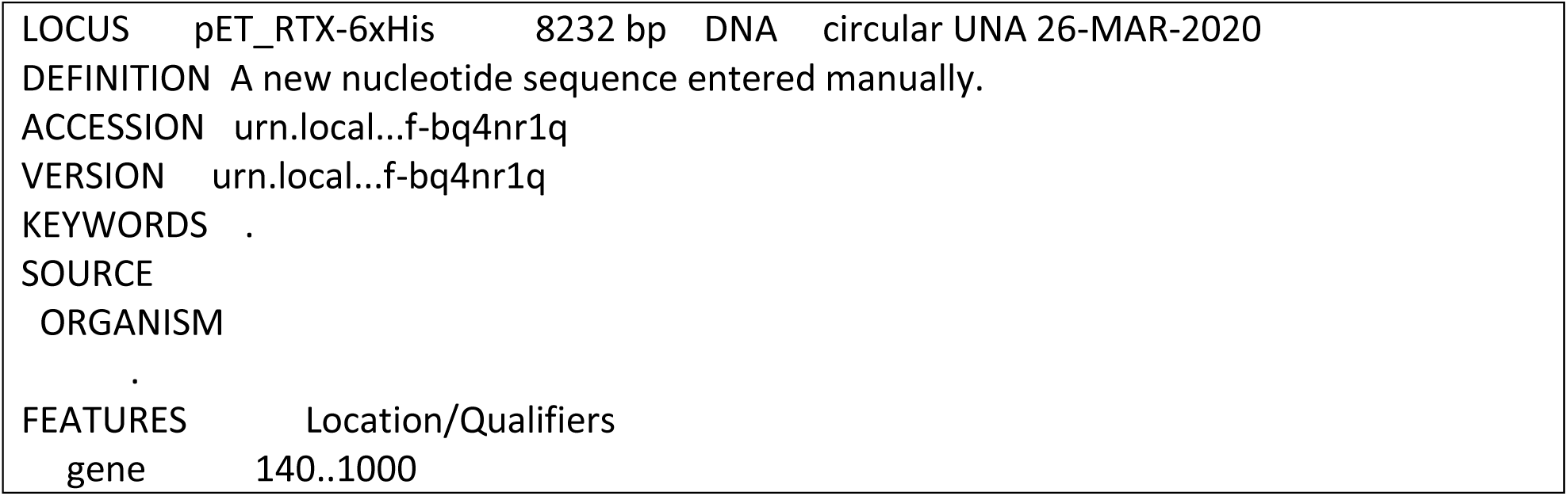
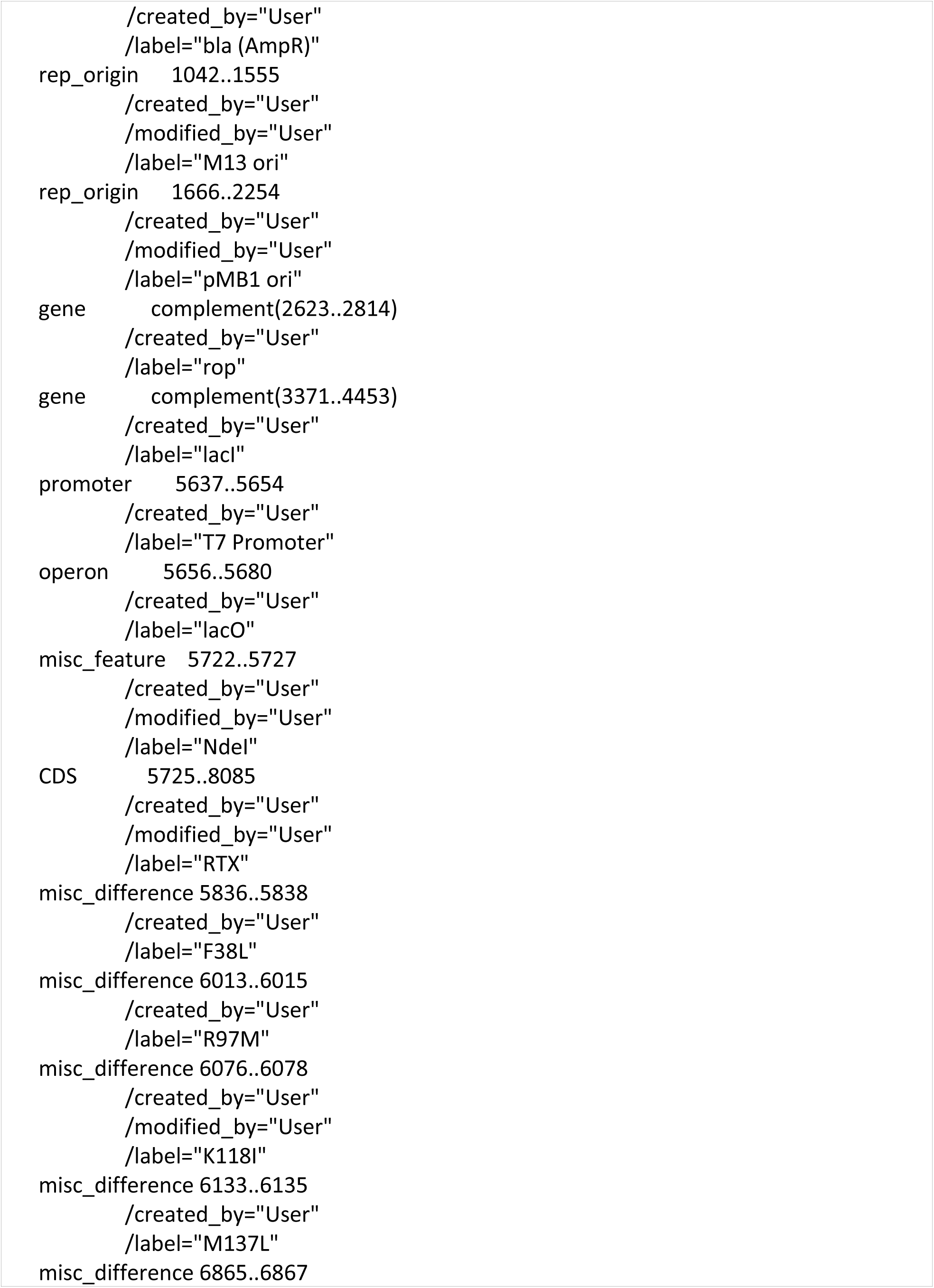
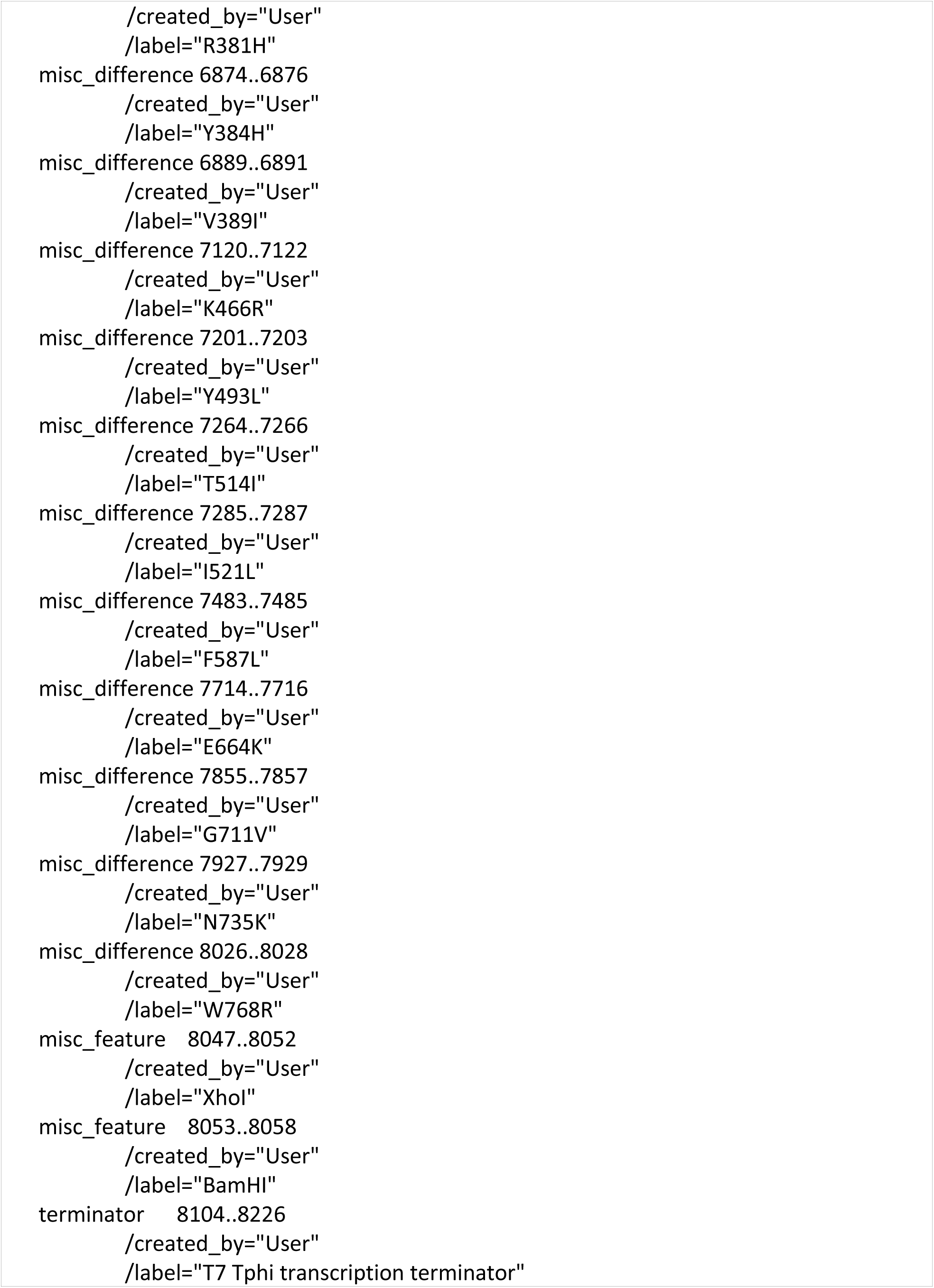
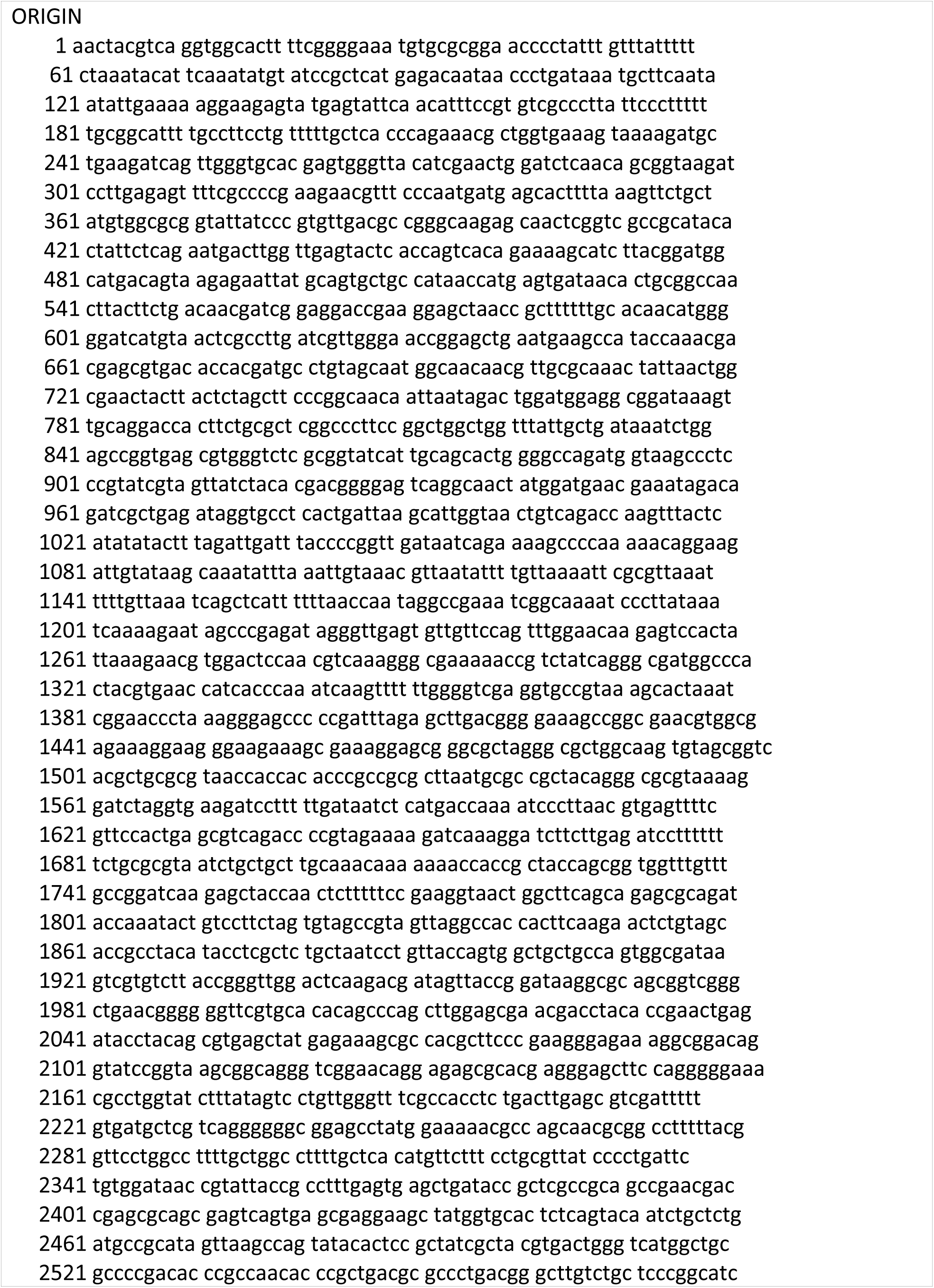
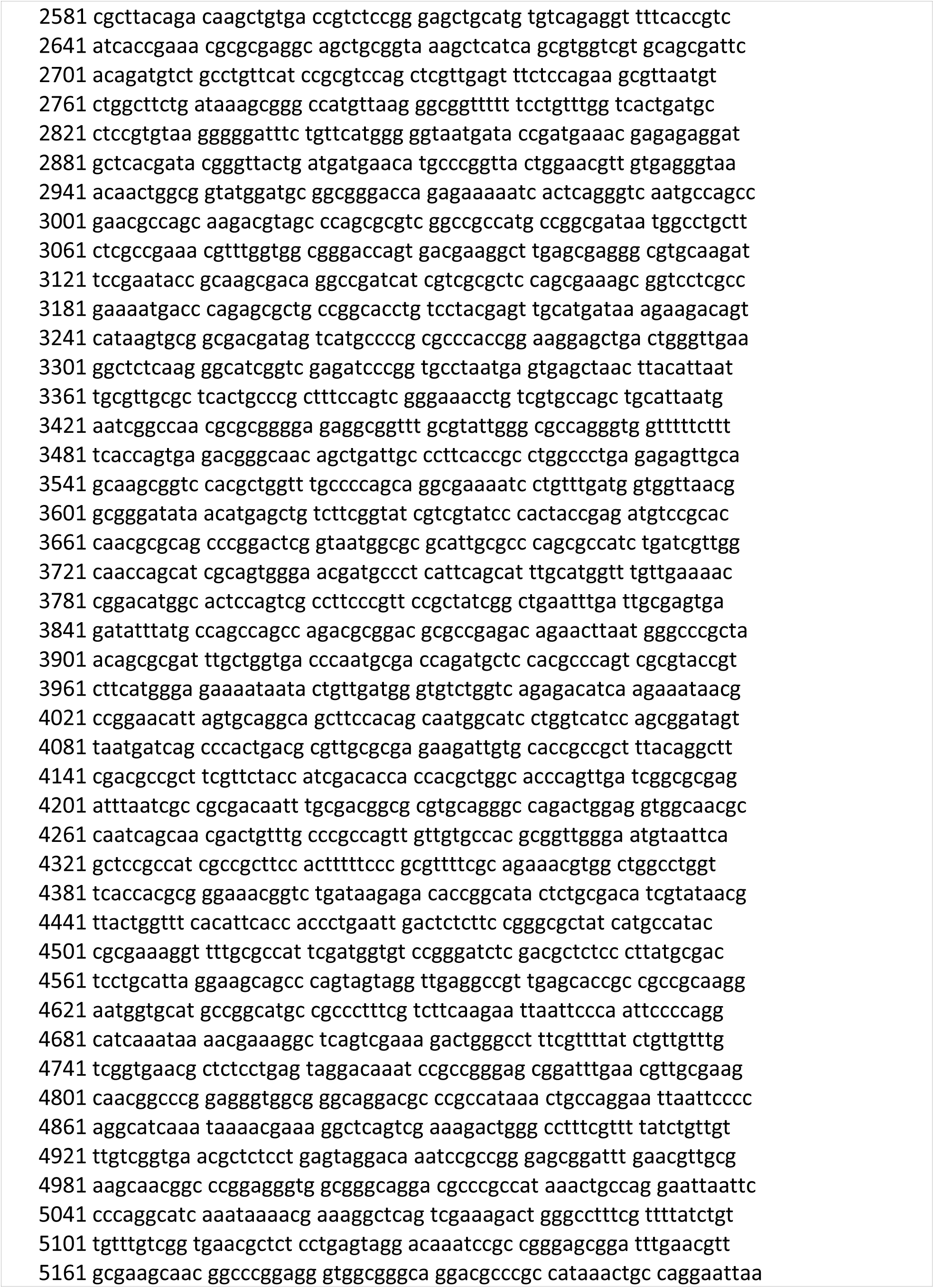
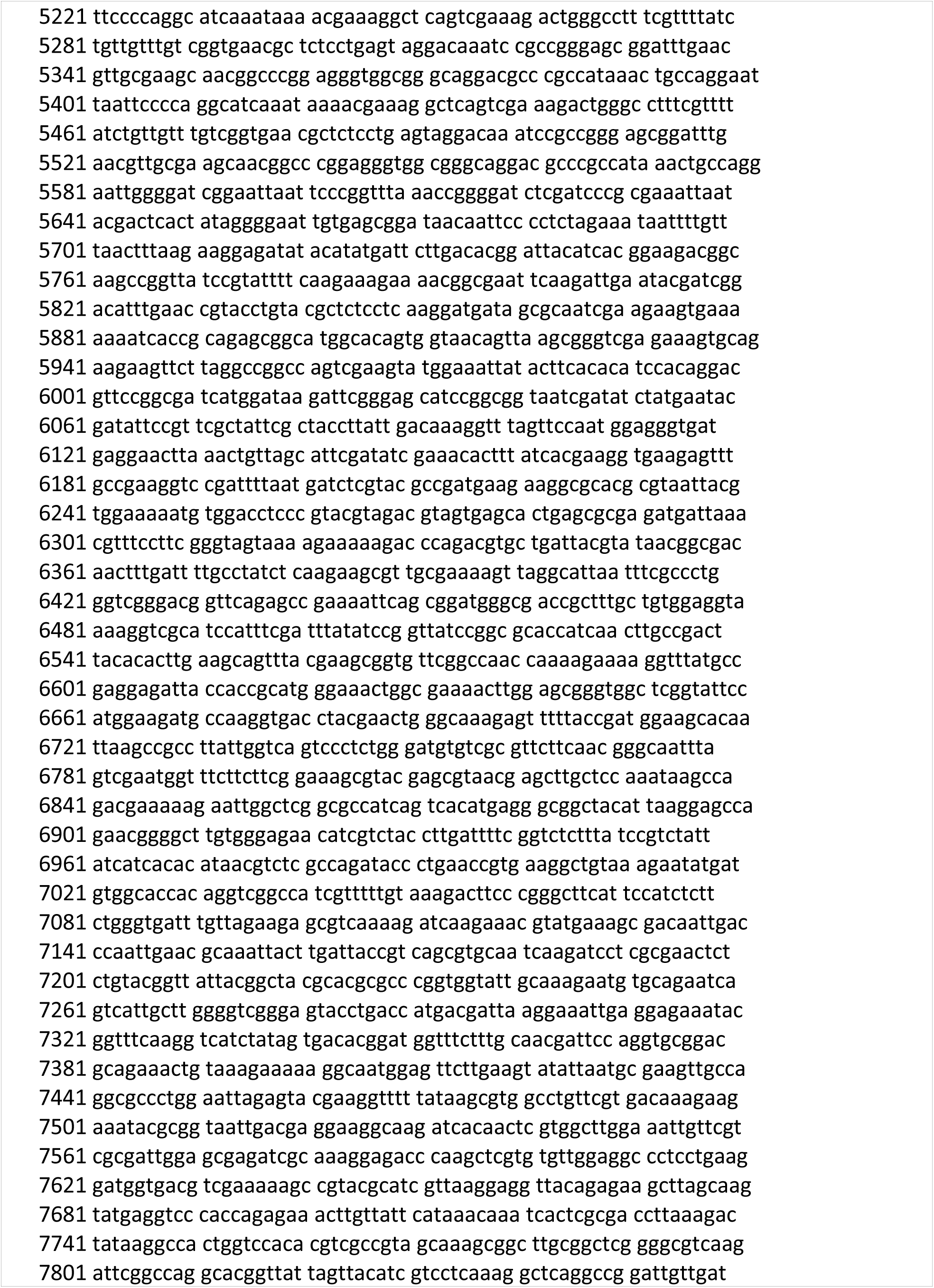
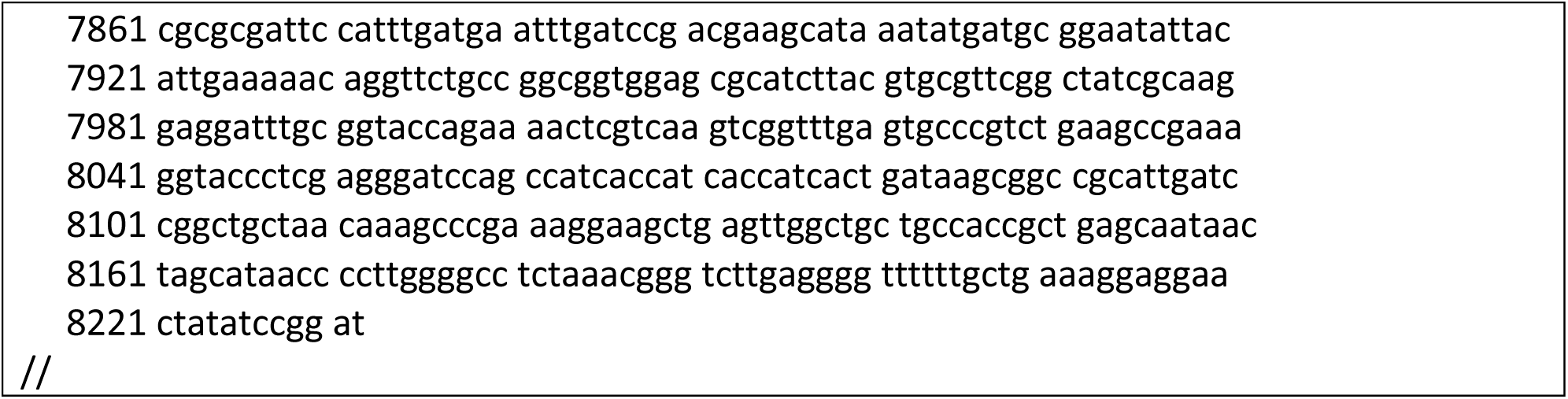
GenBank format of pET_RTX-6xHis

**Figure S2.**
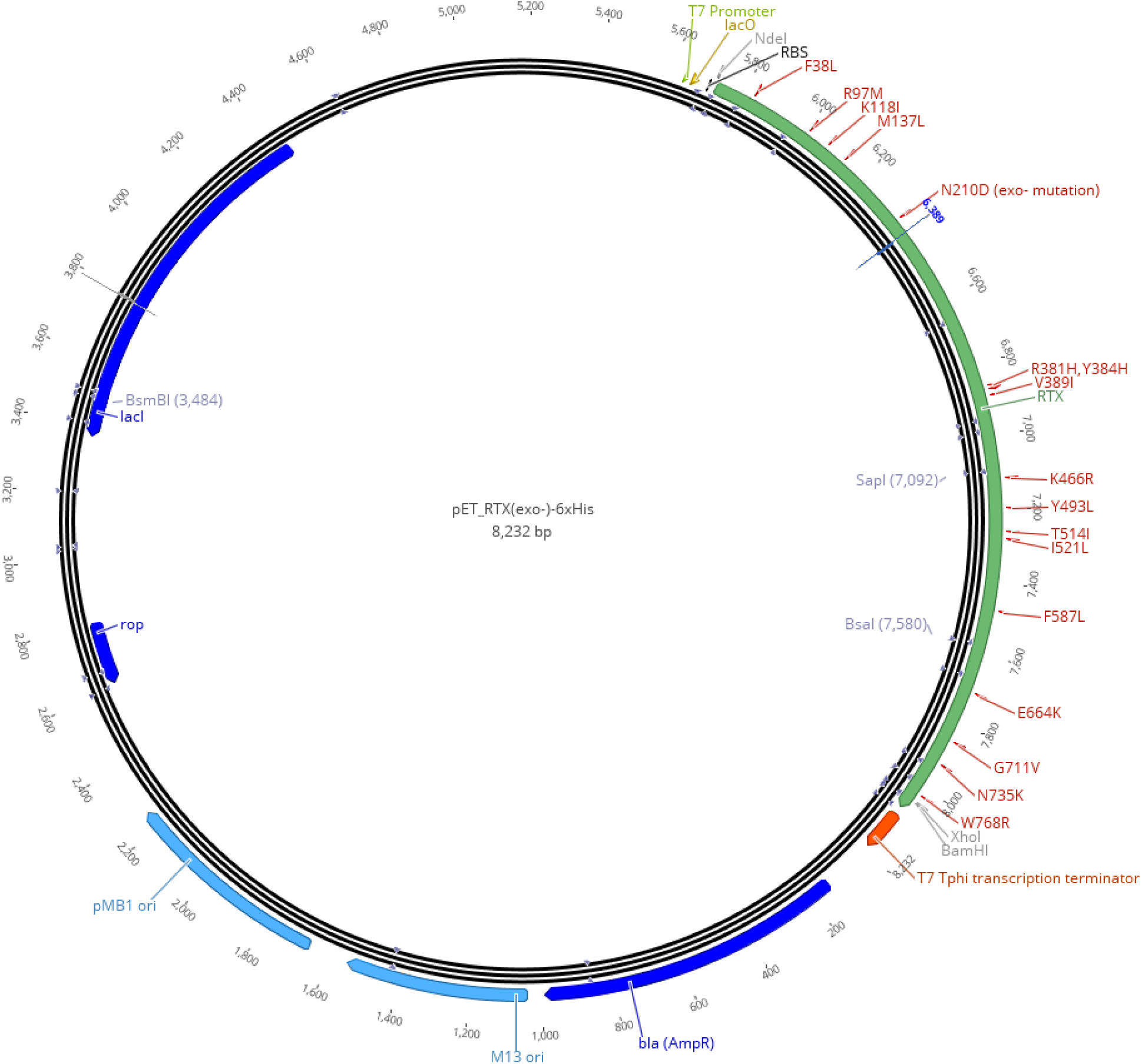
Plasmid map pET_RTX(exo-)-6xHis

**Table S2.**
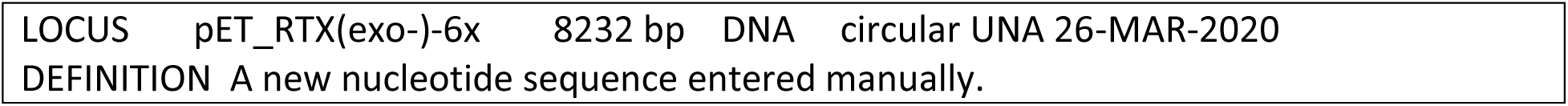
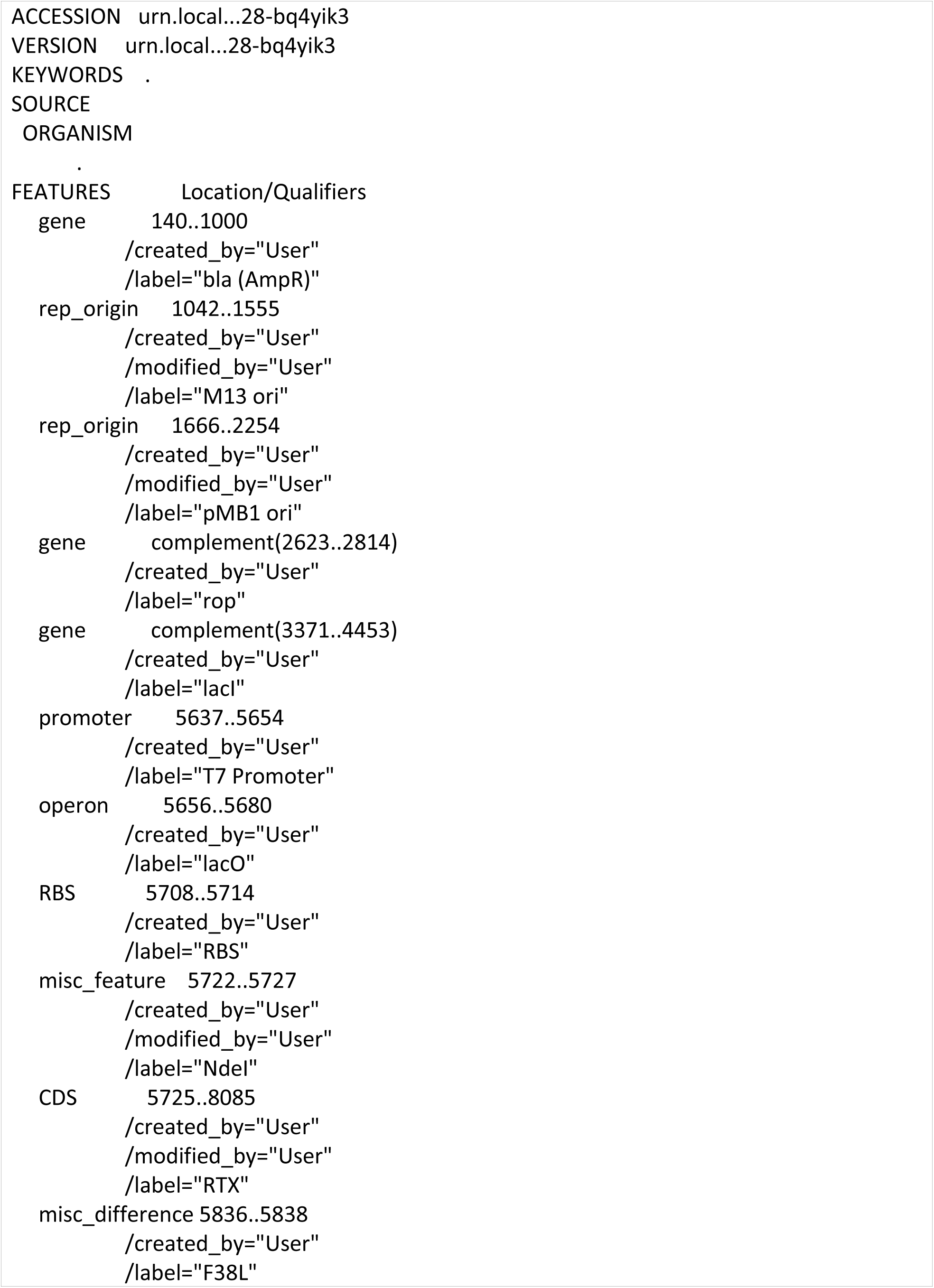
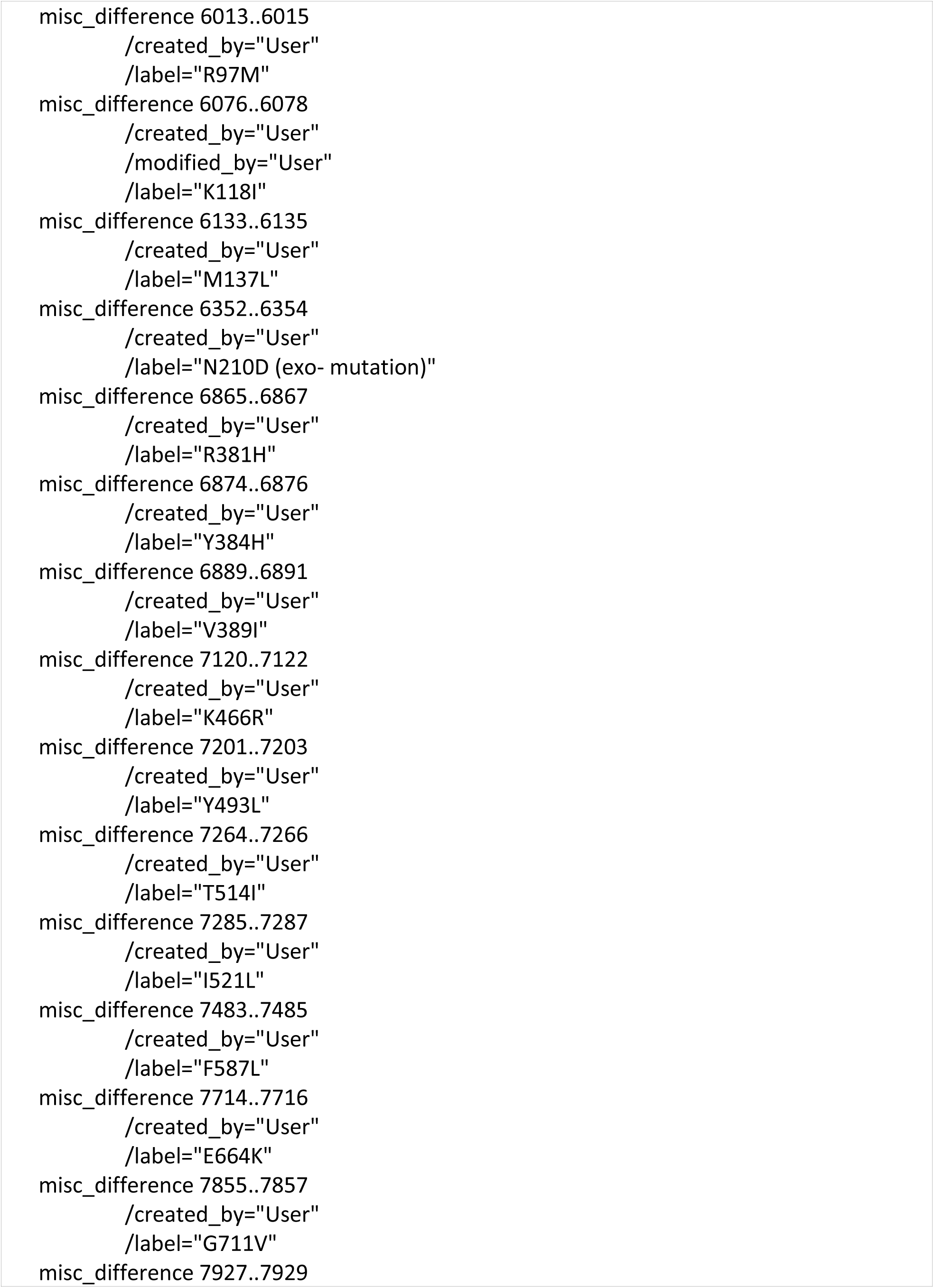
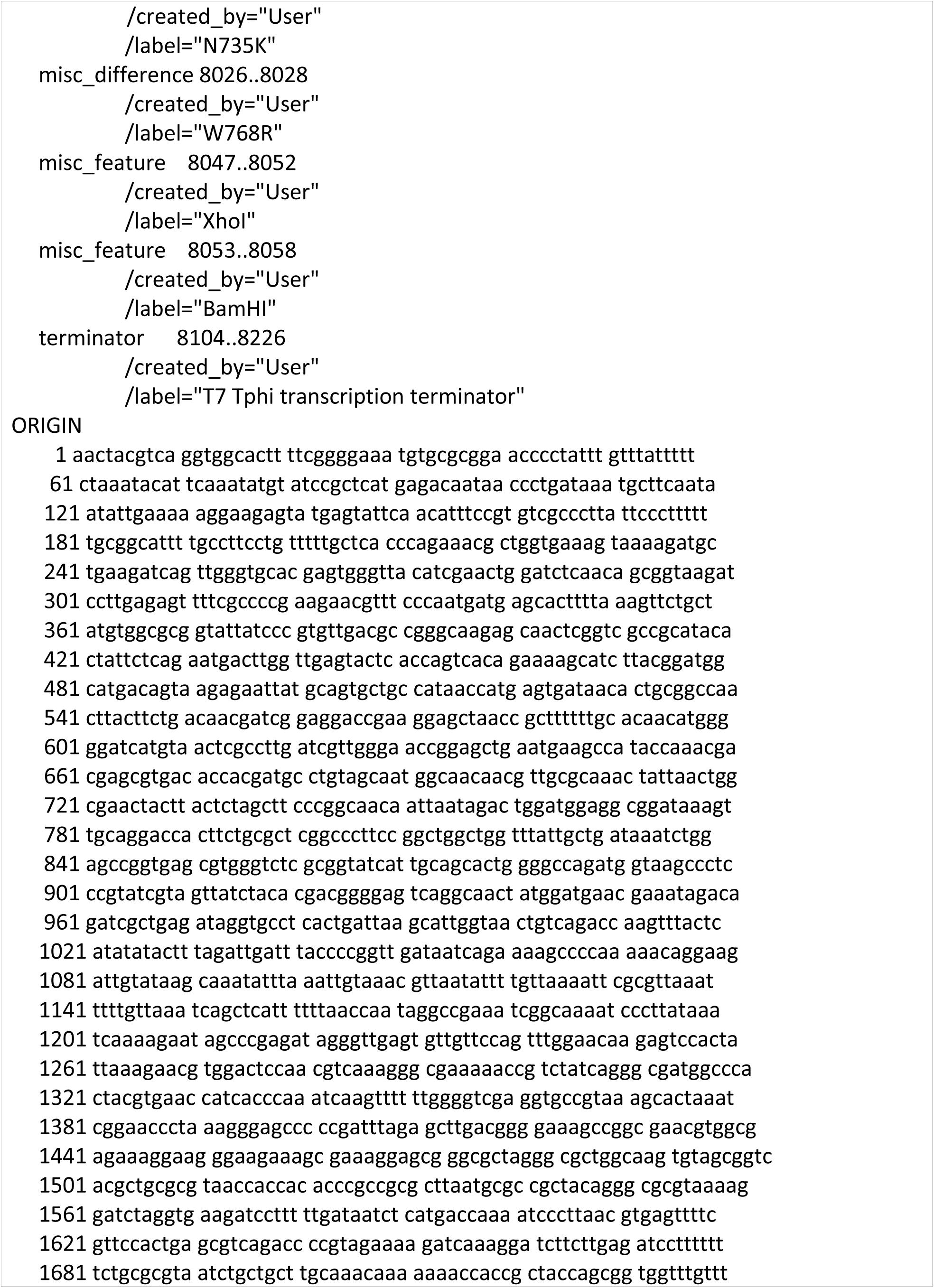
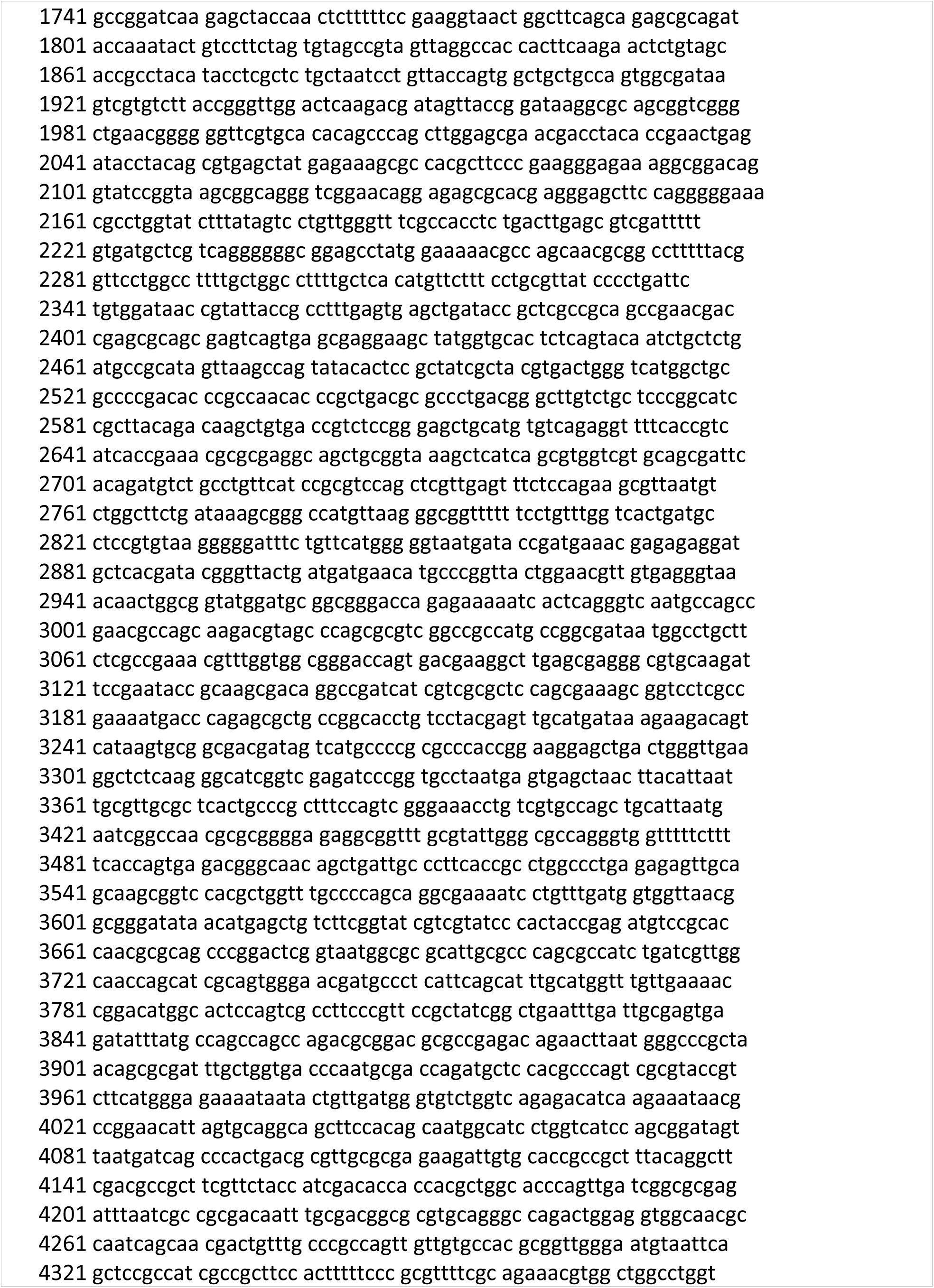
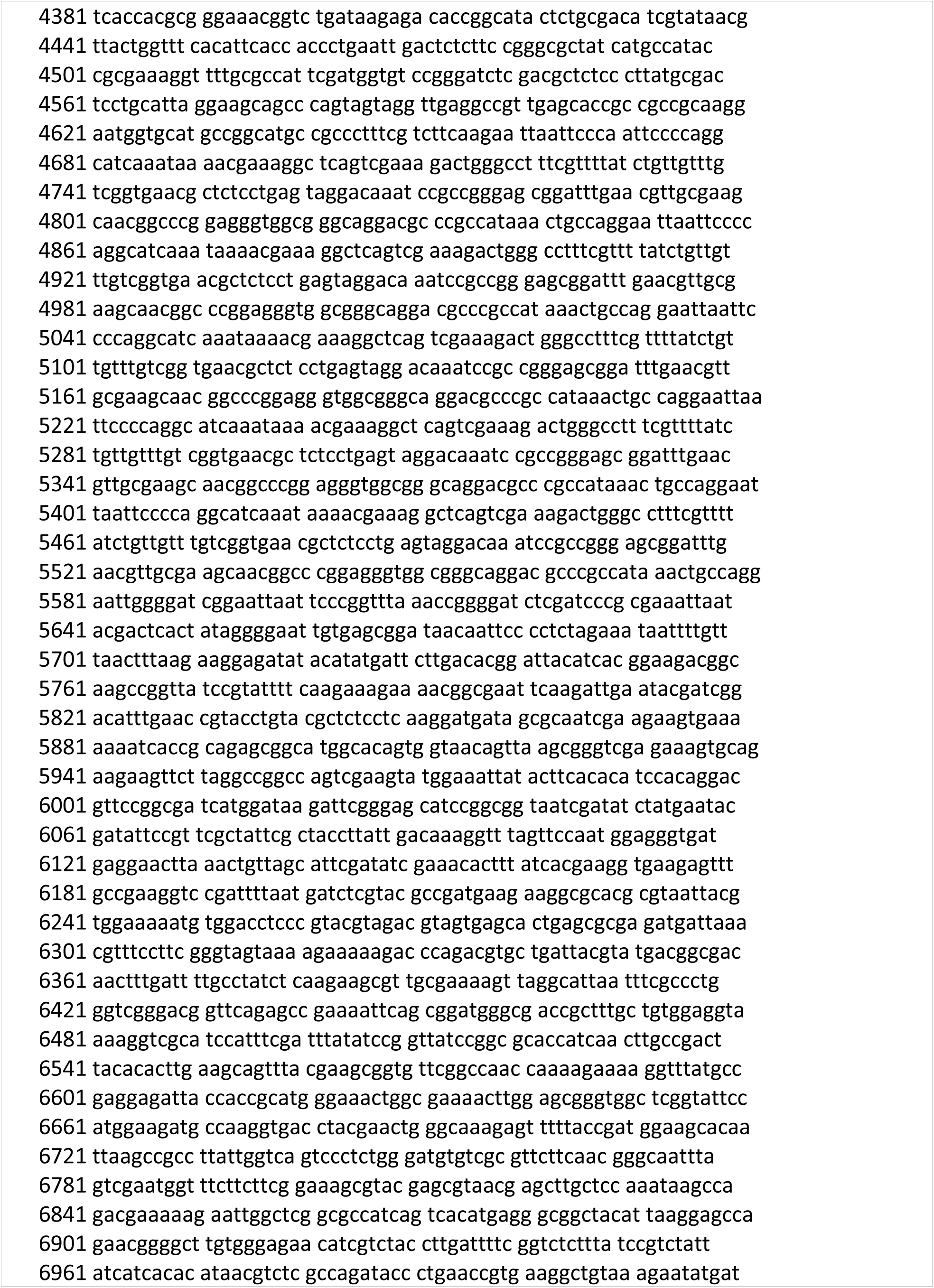
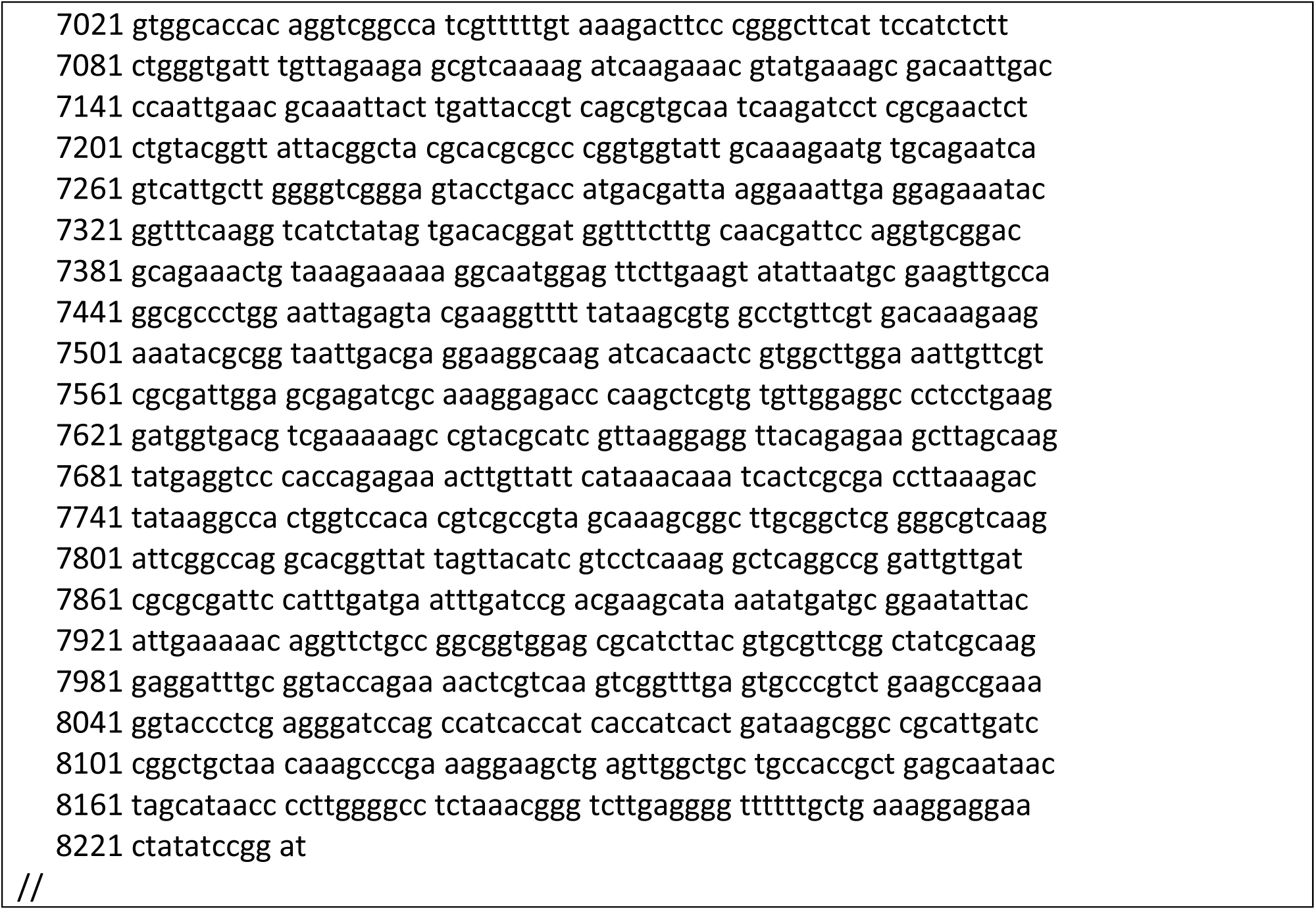
GenBank format of pET_RTX(exo-)-6xHis

